# Bayesian Workflow for Generative Modeling in Computational Psychiatry

**DOI:** 10.1101/2024.02.19.581001

**Authors:** Alexander J. Hess, Sandra Iglesias, Laura Köchli, Stephanie Marino, Matthias Müller-Schrader, Lionel Rigoux, Christoph Mathys, Olivia K. Harrison, Jakob Heinzle, Stefan Frässle, Klaas Enno Stephan

**Affiliations:** Translational Neuromodeling Unit, Institute for Biomedical Engineering, University of Zurich and ETH Zurich, Zurich, Switzerland; Max Planck Institute for Metabolism Research, Cologne, Germany; Interacting Minds Centre, Aarhus University, Aarhus, Denmark; Department of Psychology, University of Otago, Dunedin, New Zealand

**Keywords:** Translational Neuromodeling, Computational Psychiatry, Bayesian Workflow, Hierarchical Gaussian Filter (HGF), multimodal response models, robust inference

## Abstract

Computational (generative) modelling of behaviour has considerable potential for clinical applications. In order to unlock the potential of generative models, reliable statistical inference is crucial. For this, Bayesian workflow has been suggested which, however, has rarely been applied in Translational Neuromodeling and Computational Psychiatry (TN/CP) so far. Here, we present a worked example of Bayesian workflow in the context of a typical application scenario for TN/CP.

This application example uses Hierarchical Gaussian Filter (HGF) models, a family of computational models for hierarchical Bayesian belief updating. When equipped with a suitable response model, HGF models can be fit to behavioural data from cognitive tasks; these data frequently consist of binary responses and are typically univariate. This poses challenges for statistical inference due to the limited information contained in such data. We present a novel set of response models that allow for simultaneous inference from multivariate (here: two) behavioural data types. Using both simulations and empirical data from a speed-incentivised associative reward learning (SPIRL) task, we show that harnessing information from two different data streams (binary responses and continuous response times) improves the accuracy of inference (specifically, identifiability of parameters and models). Moreover, we find a linear relationship between log-transformed response times in the SPIRL task and participants’ uncertainty about the outcome.

Our analysis illustrates the benefits of Bayesian workflow for a typical use case in TN/CP. We argue that adopting Bayesian workflow for generative modelling helps increase the transparency and robustness of results, which in turn is of fundamental importance for the long-term success of TN/CP.

## Introduction

Psychiatry suffers from a dearth of tests that are based on biological or cognitive mechanisms (Kapur et al., 2012). In response, computational approaches to psychiatry begun gaining attention a decade ago (Montague et al., 2012). A particular focus has been on generative models and their potential for inference on individual disease mechanisms, as a basis for overcoming the limitations of contemporary symptom-based diagnostic classifications (Stephan & Mathys, 2014).

Computational approaches to psychiatry encompass two main branches: Translational Neuromodeling (TN) which is concerned with the development and validation of computational assays – i.e. generative models for inferring mechanisms underlying neurophysiology, behaviour, and cognition – and Computational Psychiatry (CP) which focuses on the application of these models to clinical questions such as differential diagnosis, stratification, and treatment prediction. Generative models represent a cornerstone of TN/CP because they (i) exploit the advantages of Bayesian approaches to inference, (ii) enforce mechanistic thinking, and (iii) provide estimates of system states and/or parameters that enable interpretable out-of-sample predictions by machine learning (an approach called generative embedding; for review, see Stephan et al., 2017).

However, there are numerous practical challenges for generative modelling. These include – but are not limited to – the choice of sensible priors for model parameters, identifiability both at the level of parameters and models, validation of the inference algorithm, as well as questions regarding model evaluation. Successfully managing these challenges is essential in order to obtain robust statistical results from Bayesian data analysis (BDA), which in turn is paramount to the success of TN/CP.

The motivation for this paper is twofold: First, we present a novel generative model in the framework of the Hierarchical Gaussian Filter (HGF; Mathys et al., 2011, 2014), a computational model for hierarchical Bayesian belief updating that has seen numerous applications in TN/CP (e.g. Hein et al., 2021; Iglesias et al., 2013; Lawson et al., 2017, 2021; Marshall et al., 2016; Powers et al., 2017; Sapey-Triomphe et al., 2023; Sporn et al., 2020). Our new generative model exploits two sources of information from behavioural responses, namely trial-wise predictions (binary responses) and associated response times (RTs). By exploiting two coupled streams of information for model inversion, we hoped to increase both parameter and model identifiability – issues which have proven challenging for some HGF applications (Bröker et al., 2018), particularly with binary response data (e.g. see Harrison et al., 2021; Iglesias et al., 2021). In order to acquire suitable data for this endeavour, we developed a novel speed-incentivised associative reward learning (SPIRL) task. In combination with a set of custom-built combined response models in the HGF framework, we demonstrate the utility of our dual-stream generative model, using both simulations and empirical data from the SPIRL task.

Second, we provide a worked example of *Bayesian workflow* that may usefully guide application of generative models in TN/CP, beyond the particular examples studied in this paper. This example extends previous tutorials that discussed a workflow for modelling behavioural data, but were restricted to frequentist (maximum likelihood) estimation (Wilson & Collins, 2019). We emphasise that the Bayesian workflow presented in this paper was not invented by us. Instead, it was derived from earlier proposals by others (Betancourt, 2020; Gelman et al., 2020; Schad et al., 2020; van de Schoot et al., 2021) and enriched with additional components, e.g. Bayesian comparison of model families (Penny et al., 2010). We focused on those steps of BDA that – independent from the chosen inference scheme – are particularly relevant for robustness of results from generative models.

## Methods

The analysis methods of this study were specified in a preregistered analysis plan (see section below for details). For consistency, we reuse text from our analysis plan in this Methods section, in adapted and extended form. We start by describing the behavioural learning task which was developed for this study. In what follows, we give a detailed summary of our modelling approach, both the development of novel response models combining different data modalities in the framework of the HGF as well as their application within Bayesian workflow. Our analysis had the following two central aims:

Aim 1: Provide a quantitative comparison of response models (for the HGF) that utilise binary and continuous-valued response data in different ways.

Aim 2: Provide a quantitative assessment of whether (and how) parameters characterizing subject-specific learning behaviour are associated with an individual’s measured response times.

## Analysis Plan, Data and Code Availability

A version-controlled and time-stamped analysis plan was created, detailing the analysis pipeline ex ante. The analysis plan provides a more in-depth description of the analysis protocol and is provided at https://doi.org/10.5281/zenodo.10669944. For the analysis, a custom-built pipeline was implemented in MATLAB R2019b (The MathWorks, Natick, MA, USA; code available at https://gitlab.ethz.ch/tnu/code/hessetal_spirl_analysis). Various open-source software packages were used for the analysis such as the HGF Toolbox (v7.1) as part of the ‘Translational Algorithms for Psychiatry-Advancing Science’ ([TAPAS](https://translationalneuromodeling.github.io/tapas/) v6.0.1, commit 604c568) package (Frässle et al., 2021), the Variational Bayesian Analysis Toolbox ([VBA](https://mbb-team.github.io/VBA-toolbox/), commit aa46573; Daunizeau et al., 2014) and the <u>RainCloudPlot</u> library (commit d5085be, Allen et al., 2021). Note that we are using an updated version of TAPAS compared to what was stated in the analysis plan; this version already includes functionalities to use combined response models with the HGF. All of these packages are included as submodules in the analysis code repository. The entire analysis pipeline underwent an internal code review (by a researcher not involved in the initial data analysis) in order to identify errors and ensure reproducibility of results. The data set used for the analysis is available on Zenodo (https://doi.org/10.5281/zenodo.10663643) in a form adhering to the FAIR (Findable, Accessible, Interoperable, and Re-usable) data principles (Wilkinson et al., 2016). We used Psychtoolbox-3 (Kleiner et al., 2007) to program the task of this study. The code that we used to run the experiment in the lab is available at https://gitlab.ethz.ch/tnu/code/hessetal_spirl_task.

## Behavioural Study Procedure

### Participants

In total, 91 right-handed healthy individuals (59 females, 32 males; age 24.7±4.3) completed the study. The data set consists of a pilot data set (*N* = 23) and a main data set (*N* = 68). For lack of a better term, we refer to the former as a ‘pilot’ data set; however, we emphasise that from the beginning, the designated purpose of this data set was to inform the specification of priors by independent data. All participants gave written informed consent prior to data acquisition and were financially reimbursed for their participation. The study was approved by the Ethics Commission of ETH Zurich (ETH-EK-Nr. 2021-N-05).

Our study applied the following exclusion criteria for participation: known psychiatric or neurological diseases (past or present), regular intake of medication (except contraceptives), current participation in other studies using pharmacological interventions or stimulation of brain nerves, and alcohol or drug intake during three days prior to the measurement.

Additionally, we excluded measured data sets from analysis according to quality criteria that had been pre-specified in the analysis plan. These criteria included: failure to complete the task, >10 ignored (no response and feedback) or irregular (RT <0.1s) trials, and <65% correct responses (adjusted for the probabilistic nature of the task). Eleven of the participants were excluded because they did not meet our criteria for adequate data quality and one participant was excluded due to contradicting information regarding the exclusion criteria, leaving us with a data set of *N*_*total*_ = 79 (pilot data set *N*_*pilot*_ = 20, main data set *N*_*main*_= 59; 52 females, 27 males; age 24.7±4.4).

### Behavioural paradigm

Each participant attended one experimental session during which they performed the SPIRL task (Figure 1). In this task, participants were required to learn the probabilistic association between two fractals and a monetary reward over a period of 160 trials. In each trial, participants were asked to select one of the two displayed fractals during a response window of 1.7s. After 1.7s from the trial onset, the outcome of the trial was revealed to the participant, i.e. whether the selected fractal was associated with a monetary reward on the given trial. Subsequently, a new response window started, and participants were again able to choose between the two fractals. Participants received visual feedback about their response times on every trial via a time bar. A customised payoff structure served to incentivise fast responses while still emphasizing the importance of correct predictions (Heitz, 2014). For details regarding the reward calculation, please refer to the analysis plan (Appendix A). Participants were informed about the payoff structure before the experiment. The trial structure is visualised in Figure 1a.

**Figure 1.**
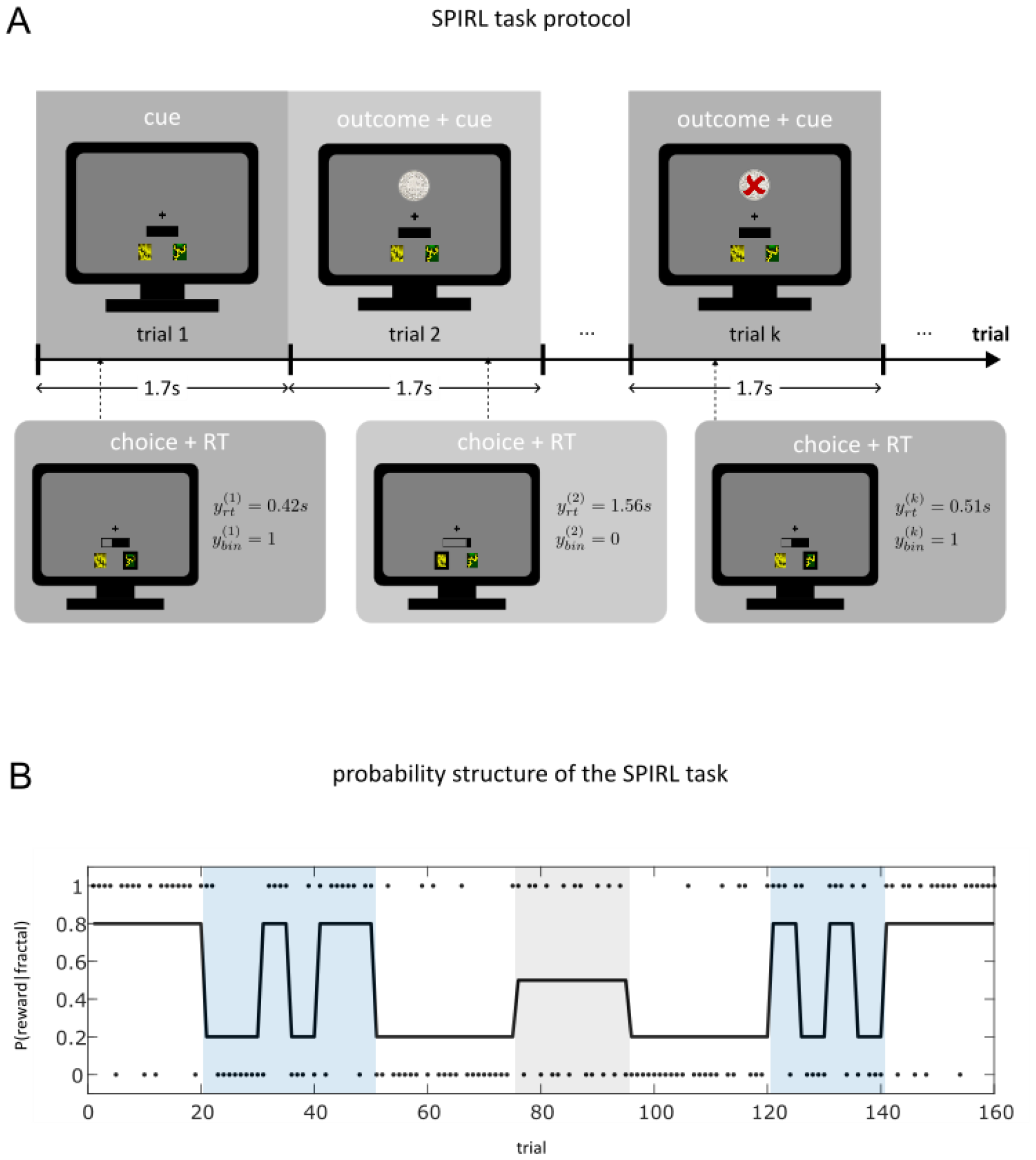
The speed-incentivised reward learning (SPIRL) task. **A** shows the trial structure of the SPIRL task protocol. A yellow and a green fractal were presented on every trial together with a time bar indicating the remaining time of the 1.7s long response window. The participants had to predict on each trial, which fractal would be rewarded monetarily. After the response window, the trial outcome was revealed (reward/no reward) concurrently with the start of the new response window of the next trial. In **B**, the probability of reward for one of the two fractals over the entire 160 trials is displayed (black line). The individual trial outcomes for this fractal are indicated by black dots (1=reward, 0=no reward). The reward probabilities of the two fractals were complementary (summing to 1) across the entire task. The colour shadings represent different phases during the task (white: stable; blue: high volatility, grey: highly unpredictable).

The probability of reward for one of the two fractals in the SPIRL task is shown in Figure 1b. The black dots indicate whether the respective fractal was rewarded on a given trial (1=reward, 0=no reward). The reward probabilities of the two fractals were designed to be complementary (summing to 1 at any given point during the experiment). Thus, the reward probability of the second fractal was simply the mirrored trajectory of the displayed trajectory, and on every trial exactly one of the two fractals was rewarded. Critically, the underlying probabilistic associations were changing over time during the experiment. The probability sequence was designed to incorporate different phases during the task indicated by colour shadings: phases with high volatility (blue), i.e. rapid switches in probability; phases where the probability was stable over a prolonged period of time (white); a phase with high uncertainty, i.e. where the outcome is unpredictable (grey). The probability sequence was fixed across participants to ensure comparability of the induced learning process. The factors fractal position (which of the two fractals was presented left) and fractal reward probability (which fractal was associated with a high reward probability in the beginning of the task) were counterbalanced across participants in the sample. Participants were told that on each trial, one of the two fractals would be rewarded and were informed about the probabilistic nature of the association between the two fractals and the monetary reward. They were also informed about the reward probabilities of the two fractals being complementary and that these probabilities were subject to changes throughout the experiment. Importantly, a priori they had no information about the values these probabilities could take as well as the order and duration of different blocks with constant probabilities.

## Analysis

### Model-agnostic analyses

The acquired behavioural data from the SPIRL task (binary responses and continuous response times) were subject to several descriptive analysis steps, mostly in the form of different visualisations of the data set. The goal of these steps was to perform a set of basic sanity checks and to identify particular characteristics of the data set. For the binary response data, adjusted correctness of the participants’ predictions (adjusted in the sense that we account for the probabilistic structure of our experiment as in (Iglesias et al., 2021), meaning that out of the total 160 trials, 122 correct predictions amount to an adjusted correctness of 100% in this task) was calculated as part of the inclusion criteria for the analysis. The descriptive analysis of the response times included different visualisations of the log-transformed response time trajectories as well as their empirical distribution in a histogram. Furthermore, we compared log RTs by task phase (stable, volatile and unpredictable according to the colour shadings in Figure 1). A one-way ANOVA of average log RTs over subjects including the factor phase was conducted as a quantitative assessment of the effect of task phase on log RTs.

### Computational modelling

For the model-based analysis, we follow the general steps of Bayesian workflow outlined in previous work (Betancourt, 2020; Gelman et al., 2020; Schad et al., 2020; van de Schoot et al., 2021) and adjusted it to our specific application at hand. Our Bayesian workflow includes the specification of an initial model space, the choice of suitable prior configurations, the choice of model inversion technique and its validation, model inversion given the empirical data set, model comparison (hypothesis testing), and model evaluation. The general steps of Bayesian workflow as well as their implementation in the context of our application are visualised in Figure 2 and described in more detail below.

**Figure 2.**
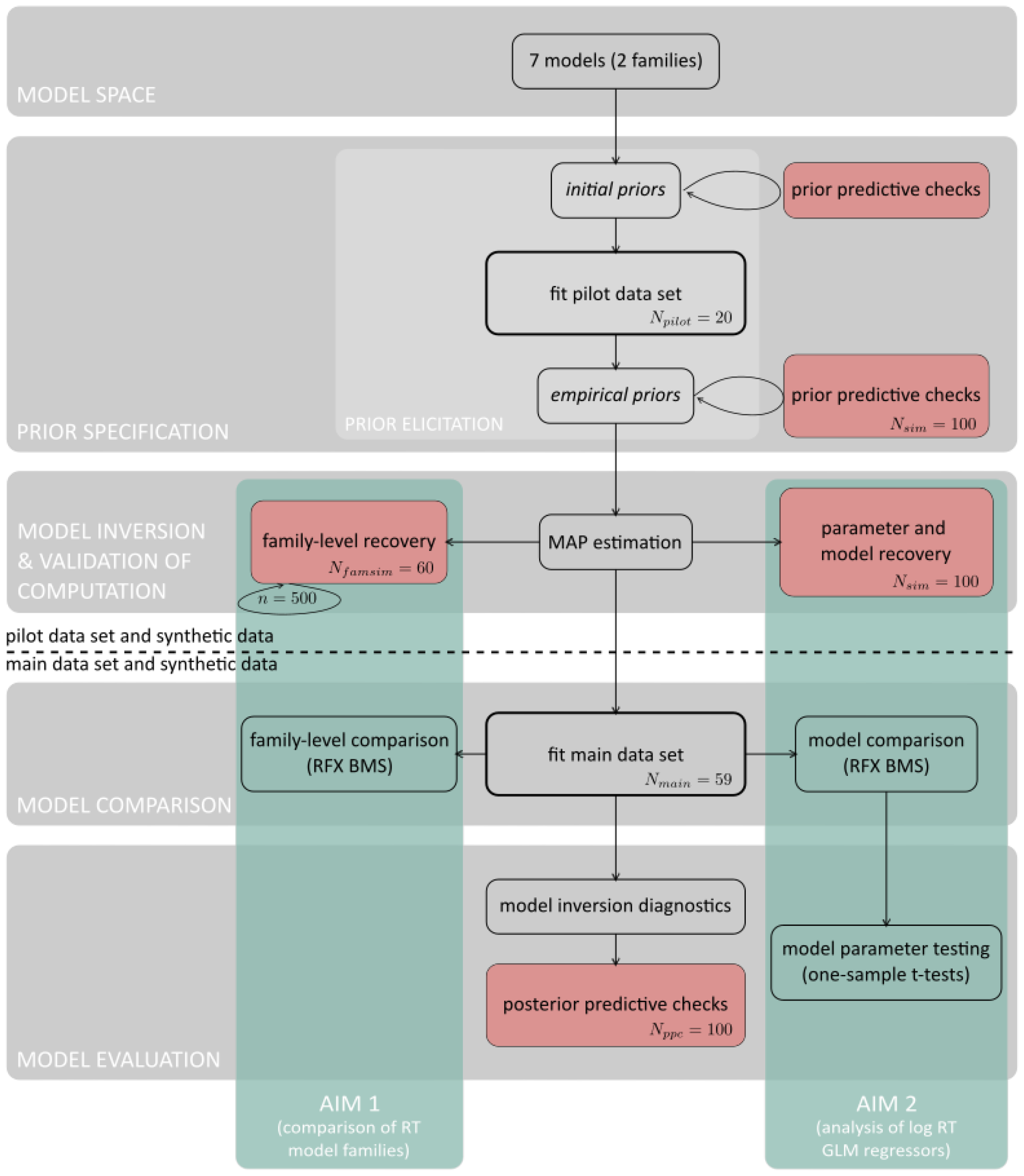
Bayesian workflow for generative modelling in Computational Psychiatry. The general steps of Bayesian workflow are visualised. These include the specification of a model space, prior specification, model inversion and validation of computation, model comparison as well as model evaluation. For each step, the concrete implementation in the context of our application is shown. Above the dashed line are analysis steps that involve the pilot data set as well as synthetic data generated using parameter values sampled from the priors. Below the dashed line are analysis steps including the main data set and synthetic data generated using parameter values sampled from the posteriors. Filled red boxes are analysis steps involving synthetic data. The green boxes highlight analysis steps that specifically refer to our research aim 1 (comparison of RT model families) and research aim 2 (assessment of individual RT model parameters).

### Model space

The set of models was based on hierarchical Gaussian filtering (HGF), a generative modelling framework for hierarchical Bayesian belief updating, describing the evolution of (hidden) states and how these give rise to the sensory inputs (*u*) an agent receives (Mathys et al., 2011, 2014). We used a 3-level enhanced HGF (eHGF) for binary inputs for the current application, where the states evolve as Gaussian random walks (GRW) at all but the first level and the step-size of the GRW at any given level depends on the next higher level (variance-coupling). The probability of the binary states at the lowest level originates from a sigmoid transformation of the quantity at the second level. The generative model of the eHGF is visualized as graphical model in the upper part of Figure 3 and described by the following set of equations

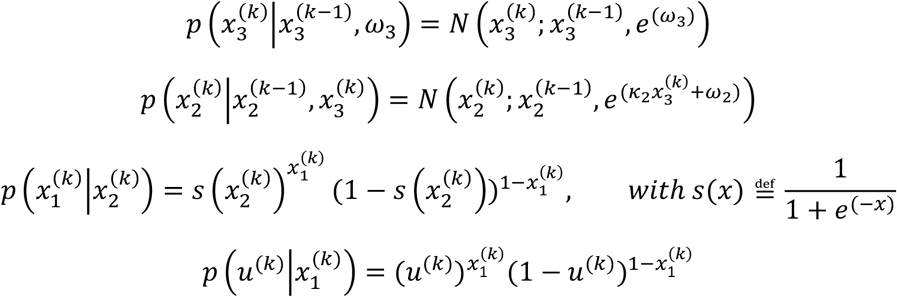

Further details and equations of the generative model can be found in Mathys et al. (2011).

**Figure 3.**
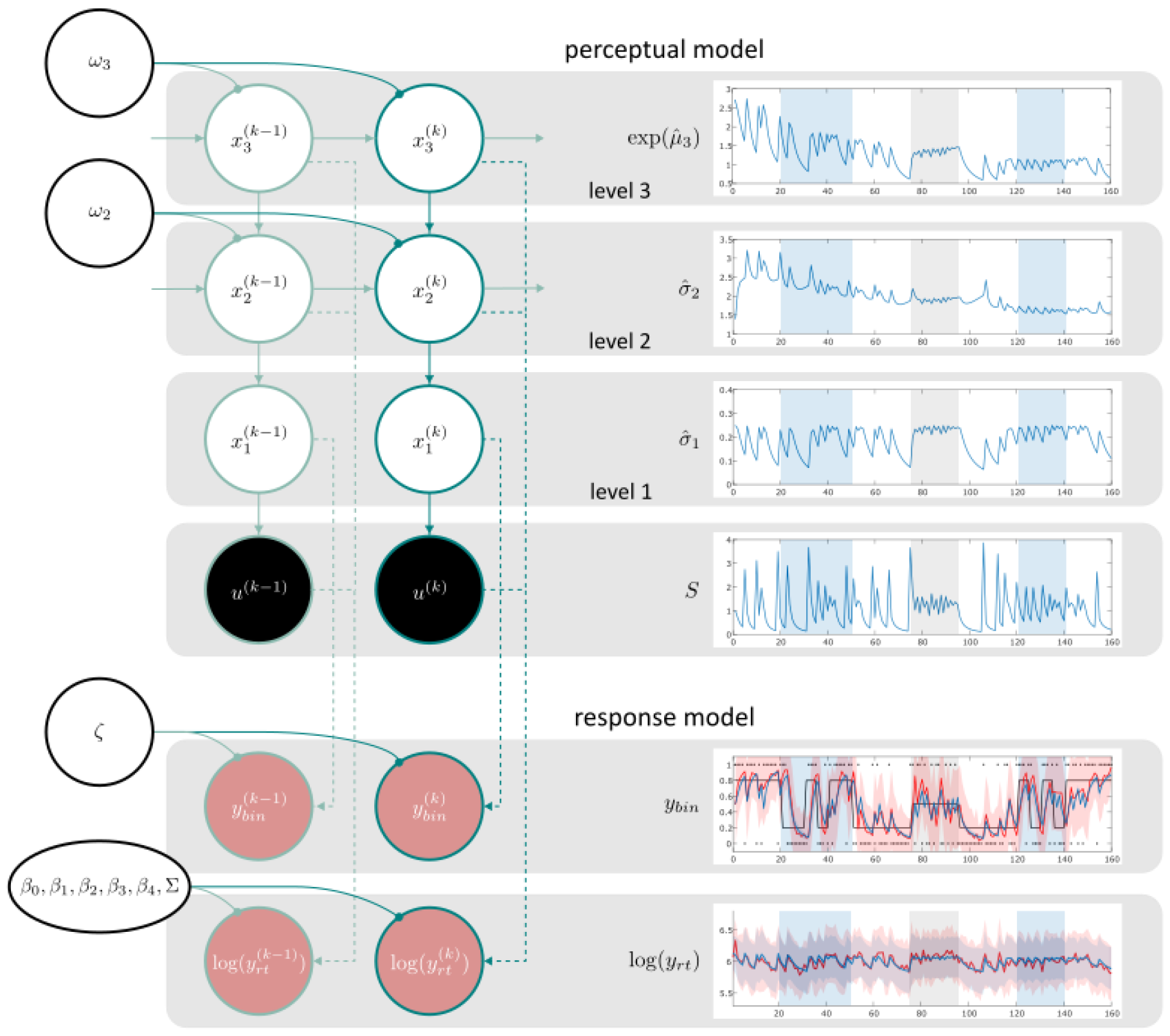
Graphical model representation of M1. In the left part of the figure, a schematic representation of the generative model of the 3-level eHGF for binary inputs (perceptual model) is presented above the dashed arrows. Below the dashed arrows, the response data modalities are visualised (response model). Shaded circles represent known quantities (inputs shaded black, response data shaded red). Unshaded circles represent estimated time-independent parameters (black circles) and time-varying states with trial indices as superscript. Dashed lines indicate the result of an inferential process, i.e. the response model builds on a perceptual model inference. Solid lines indicate generative processes. Dark turquoise lines indicate the probabilistic network on trial *k*. Light turquoise lines indicate the network at other points in time. On the right side of the figure, average belief trajectories of the perceptual model are shown in blue in the upper four panels. The four belief trajectories represent the average trajectory over participants (*N*_*main*_= 59) of the states that enter the log RT GLM of M1. In the lower two panels, average response data over participants are shown. For the binary response modality, the red line represents average binary prediction 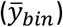, the blue line represents the average belief about the probability of the outcome 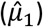 according to M1. For the continuous response modality, the red line represents average log-transformed RTs and the average predicted log RTs by M1 are shown in blue.

The 3-level eHGF for binary inputs represents a concrete implementation of the meta-Bayesian ‘observing the observer’ framework, where an observation or response model serves to specify a mapping from inferred beliefs of an agent to observed responses as recorded during our experiment (Daunizeau et al., 2010a; Daunizeau et al., 2010b). The response model uses the perceptual model indirectly via its inversion (Mathys et al., 2014). In this study, we augmented the perceptual model (eHGF) with a novel set of response models combining binary responses and continuous-valued response times.

In the eHGF, variational inversion of the agent’s model of the world (perceptual model) using Variational Bayes (VB) under a mean-field approximation and a quadratic approximation to the variational energies give rise to a set of analytical trial-by-trial update equations that resemble the general structure of Reinforcement Learning (RL) models but explicitly represent inferential uncertainty. For example, the update of the belief at the second level of the eHGF about the cue-outcome contingency takes the form of

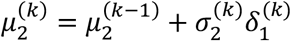

where superscripts refer to trial indices and subscripts to the level in the hierarchy of the model. In other words, the prediction 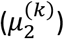 at trial *k* is equal to the sum of the prediction at the previous trial 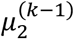 and a learning rate 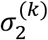 multiplied with a prediction error term 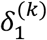. Here the prediction error 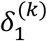 describes the mismatch between the actual (*u*^(*k*)^) and the predicted sensory input 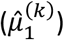

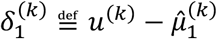

where the predicted sensory input is simply a sigmoid transformation of the belief at the second level

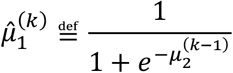

Notably, in the HGF, the learning rate 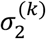 is dynamic and corresponds to a function of the temporary evolving uncertainty of the agent’s belief about the cue-outcome contingency 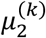. A full account of the update equations describing the evolution of beliefs at all levels of the 3-level eHGF for binary inputs can be found in the analysis plan (Appendix A3).

We defined a set of combined response models that are all paired with a 3-level eHGF for binary inputs (perceptual model). The combination of binary responses and continuous response times in these response models is accomplished by summing the log-likelihood of two individual response models, assuming independence of the two response data modalities conditional on the parameters of the perceptual model. All seven models in our model space are listed in Table 1.

**Table 1.** Model Space. All seven models in our model space are composed of a perceptual (Prc) model and an observation (Obs) or response model. The perceptual model as well as the binary part of the response model is held constant across all seven models. The equations of the log RT GLMs (continuous part of the response model) of M1-M4 (family of informed RT models) contain belief trajectories of the perceptual model as regressors. The update equations for the perceptual model (eHGF) are listed in the analysis plan. M5-M7 (uninformed RT family) predict log RTs independent of the perceptual model.

| Model | Prc model | Obs model<br>(continuous) | Obs model<br>(binary) |
| --- | --- | --- | --- |
| M1 | eHGF | $\log(y_{rt}^{(k)}) \sim N(\beta_0 + \beta_1 S^{(k-1)} + \beta_2 \hat{\sigma}_1^{(k)} + \beta_3 \hat{\sigma}_2^{(k)} + \beta_4 e^{\hat{\mu}_3^{(k)}}, \Sigma)$<br>with<br>$S^{(k-1)} \stackrel{\text{def}}{=} \begin{cases} -\log_2(\hat{\mu}_1^{(k-1)}) & \text{if } u^{(k-1)} = 1 \\ -\log_2(1 - \hat{\mu}_1^{(k-1)}) & \text{if } u^{(k-1)} = 0 \end{cases}$ | Unit-square sigmoid |
| M2 | eHGF | $\log(y_{rt}^{(k)}) \sim N(\beta_0 + \beta_1 \sigma_2^{(k)} + \beta_2 \hat{\mu}_3^{(k)}, \Sigma)$ | Unit-square sigmoid |
| M3 | eHGF | $\log(y_{rt}^{(k)}) \sim N(\beta_0 + \beta_1 \hat{\sigma}_1^{(k)} + \beta_2 \sigma_2^{(k)} + \beta_3 e^{(\kappa_2 \mu_3^{(k-1)} + \omega_2)}, \Sigma)$ | Unit-square sigmoid |
| M4 | eHGF | $\log(y_{rt}^{(k)}) \sim N(\beta_0 + \beta_1 \sigma_2^{(k)} + \beta_2 e^{(\kappa_2 \mu_3^{(k-1)} + \omega_2)}, \Sigma)$ | Unit-square sigmoid |
| M5 | eHGF | $\log(y_{rt}^{(k)}) \sim N(\beta_0, \Sigma)$ | Unit-square sigmoid |
| M6 | eHGF | $\log(y_{rt}^{(k)}) \sim N(\beta_0 + \beta_1 \frac{k}{K}, \Sigma)$ | Unit-square sigmoid |
| M7 | eHGF | $\log(y_{rt}^{(k)}) \sim N(\beta_0^{(k)}, \Sigma)$<br>with<br>$\beta_0^{(k)} \stackrel{\text{def}}{=} \begin{cases} a, & \text{if } y_{bin}^{(k-1)} = u^{(k-1)} \\ b, & \text{if } y_{bin}^{(k-1)} \neq u^{(k-1)} \end{cases}$ | Unit-square sigmoid |

As binary part of the response model, we used a unit-square sigmoid function mapping from the belief 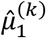 at the lowest level of the eHGF that the next outcome will be 1 onto the probabilities 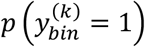 and 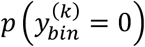 that the agent will choose response 1 or 0 (for simplicity, in the following equation we use 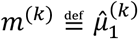 and we omit time indices on *y* and *m*):

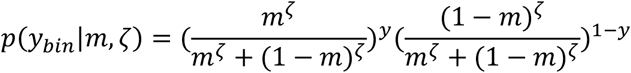

Here, the parameter *ζ*determines the steepness of the sigmoid function and is referred to as inverse decision temperature or inverse decision noise. For reasons of comparability, the binary part was kept constant across all response models in our model space.

For the continuous part of the response models, we used seven different variants of a general linear model (GLM) to predict the log-transformed RTs on a trial-by-trial basis. The regressors of models 1-4 (M1-M4) are belief trajectories from the perceptual model (eHGF), e.g., an individual’s estimated uncertainty about the outcome 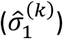 at trial *k*. This was inspired by the approaches used in previous work where reaction time data had been modelled using the HGF framework (Lawson et al., 2017, 2021; Marshall et al., 2016). A set of alternative combined response models (M5-M7) predicted RTs independently of the perceptual model (eHGF); including these models allowed us to test whether informing RTs by inferred states from the eHGF would improve the models at all.

Our model space can thus be divided into 2 families based on the continuous part of the response models (since the perceptual model and the binary part of the response models were held constant). M1-M4 are ‘informed’ RT models and M5-M7 are ‘uninformed’ RT models. Note that Bayesian model comparison is based on the (log) evidence and therefore requires the data to be the same across all models. In our case, it is thus not possible to include a model that does not model RT at all in the model comparison.

The GLM equations of M1-M7 are listed in Table 1, and a graphical representation of M1 is presented in Figure 3. Concerning the first (informed) set of models, the GLM of M1 was adapted from a model used by Lawson et al. (2017) and tailored towards the specific properties of the SPIRL task. Models M2-M4 are custom built response models with the following underlying motivation: Firstly, the goal was to reduce the number of regressors entering the GLM and hence the complexity of the models compared to M1. Secondly, the idea was to include different types of uncertainty from different levels of the hierarchy described by the perceptual model (eHGF). The eHGF accommodates various forms of uncertainty. Two sources of uncertainty, informational and environmental uncertainty, respectively, are represented in the update equations of the eHGF. M2 includes estimates of informational uncertainty 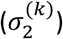 and phasic volatility 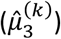. M3 includes estimates of informational uncertainty at the outcome level 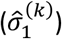 and at the second level 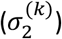 as well as environmental 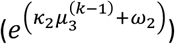 uncertainty whereas M4 only includes estimates about the two latter quantities. For a more detailed discussion of the different forms of uncertainty captured by the eHGF, please see Mathys et al. (2014).

By contrast, the second (uninformed) set of models, M5-M7, have response time GLMs where the regressors are independent of the perceptual model. M5 is a null model, representing the assumption that log RTs are simply noise around a constant intercept. M6 allows for a linear decay over time in the response times, and M7 corresponds to a null model with two different intercepts for correct and incorrect response trials (referring to the binary response data).

### Prior specification

#### Prior elicitation

Since the central questions of the present study are addressed by Bayesian model comparison (whose central quantity, the model evidence, takes into account the models’ priors), the specification of priors is a critical step. We aim to specify priors that lay in the range of actual human behaviour for the present task, while at the same time minimising the subjectivity involved in the choice of priors. In order to avoid problems of double-dipping, we made use of an independent data set, i.e. the 20 participants from our pilot study, for the elicitation of prior densities. Recently, similar approaches have been used for comparable modelling endeavours (Harrison et al., 2021; Schöbi et al., 2021). We used a two-step procedure for elicitation of prior densities that were subsequently used in the analysis of the main data set (for the results, see Supplementary Material S2). We estimated the sufficient statistics (mean and variance) of the prior densities as follows (all priors take the form of a normal distribution):

1. Inversion of M1-M7 given the pilot data set (*N*_*pilot*_ = 20) using *initial prior* means and variances. These *initial prior* means and variances represent the default prior configurations of the eHGF in the HGF Toolbox.
2. For every model *m*, we defined:
  a. a new prior mean *pE*_*m*_ as the robust mean estimate over Maximum a posteriori (MAP) estimates obtained in step 1.
  b. a new prior variance *pC*_*m*_ as the robust variance estimate over MAP estimates obtained in step 1.

For the robust estimation of mean and variance, we used a variant of the minimum covariance determinant (MCD) method, namely the FAST-MCD algorithm developed by (Rousseeuw & Van Driessen, 1999), as implemented in the ‘robustcov’ function in MATLAB R2019b. We refer to this new set of priors, that we estimated via the above described procedure, as *empirical priors* (as informed by the pilot data set). Please note that our procedure is distinct from commonly employed ‘empirical Bayesian’ procedures that estimate priors and parameter values from the same data set, using a hierarchical model. The advantage of our procedure is that the data which inform the choice of priors is fully independent from the data that inform parameter estimation.

#### Prior predictive checking

In order to get an intuition for the range of behaviours which can be generated by our models under the *initial* and *empirical priors*, we looked at their prior predictive distributions. More specifically, we randomly sampled 100 parameter values from the respective prior densities and simulated belief trajectories and responses. Adequacy of our chosen prior distributions was determined in a qualitative manner (visualisations) as well as in a quantitative fashion in the form of different recovery analyses (see paragraph ‘Validation of model inversion’). More detailed information describing our choice of priors and prior predictive checking can be found in the analysis plan.

### Model inversion

We inverted the generative models using approximate Bayesian inference as implemented in the HGF Toolbox. MAP estimates are computed as the minimum of the negative log joint using gradient-based optimisation techniques. By default, the HGF Toolbox uses an optimisation algorithm from the quasi-Newton methods family (Broyden-Fletcher-Goldfarb-Shanno algorithm; BFGS) (Broyden, 1970; Fletcher, 1970; Goldfarb, 1970; Shanno, 1970). The covariance of the posterior was obtained under a Laplace approximation to the negative log joint at the MAP. This allowed for the specification of credible intervals on the obtained parameter estimates as well as the calculation of an approximate log model evidence (LME) as a measure of ‘model goodness’. For the present analyses, a multi-start optimisation approach was used to alleviate issues with the optimisation algorithm getting stuck in local extrema of the objective function. Specifically, 400 different starting points were used for the inversion of each subject and model. Of these 400 starting points, one always corresponded to the prior mean values of the different parameters (representing the default setting in the HGF Toolbox), whereas the other 399 starting points were randomly sampled values from the respective prior density of the parameters.

It should be noted that this choice of inference technique means that we are dealing with a simplified case of BDA. That is, we only obtain a point estimate of the posterior and approximate the posterior uncertainty. An advantage of our model inversion procedure is the computational efficiency and the avoidance of concerns about convergence, as in the case of Markov Chain Monte Carlo (MCMC) procedures. Having said this, for the specific purpose of our paper, the choice of model inversion approach is not critical.

### Validation of model inversion

To examine the identifiability of parameters from the models in our model space, we performed a set of and recovery analyses using synthetic data (for a tutorial introduction to recovery analyses, albeit in the context of maximum likelihood estimation, see Wilson and Collins (2019)). Examining the identifiability (or recoverability) of parameters and models can also be seen as a test of face validity, i.e. asking whether the models actually do what they are supposed to do: allowing for veridical parameter estimates and representing a distinct explanation of observed data that can be distinguished from other explanations. This step is an important part of Bayesian workflow because it establishes a boundary between the type of questions that can be addressed using the model space at hand, and those questions for which a meaningful answer cannot be expected. In other words, we assessed whether in principle (i.e., knowing the ‘true’ data-generating models and parameter values), we would be able to identify the data-generating model as the model that explained the data best (model identifiability) and the parameter values of the data generating model (parameter recovery), respectively. In a second step, we assessed whether data-generating model families could be recovered using family-level comparisons.

For each model, we generated a synthetic data set (*N*_*sim*_ = 100) by randomly drawing 100 samples from the *empirical prior densities* and plugging these values into the likelihood function. We then fit the synthetic data sets by each of the models in our model space under the respective *empirical priors*. Parameter recovery was assessed by visually comparing simulated parameter values to MAP estimates obtained using the data-generating model and by calculating Pearson correlation coefficients *r*. Model identifiability was quantified both as the proportion of correctly identified models according to approximate LME scores in a classification analysis, and calculating protected exceedance probabilities (PXP) as part of random-effects (RFX) Bayesian model selection (BMS) (Rigoux et al., 2014; Stephan et al., 2009). Moreover, we computed a balanced accuracy score for the LME winner classification and compared it to the upper bound of a 90%-CI when assuming each model to be selected with equal probability. Family level recovery was quantified using approximate LME scores obtained during the model identifiability analysis to create 5’000 synthetic data sets of *N*_*famsim*_ = 60 subjects each, with different ratios of data generating families. We compared true family frequencies with expected posterior family frequency (Ef) and family exceedance probabilities (XP) resulting from family-level RFX BMS (Penny et al., 2010). More details on the background of BMS methods can be found in the next section (‘Model comparison’) and in the referenced literature.

### Model comparison

After careful investigation of our model space and validation of the chosen inference algorithm, we inverted the models given the main data set in order to address the following aims:

**Aim 1**. Perform a quantitative comparison of response models that utilise binary and continuous-valued response data in different ways. More specifically, we tested whether the family of informed RT models explained the collected data better than the uninformed RT model family, or vice-versa. To this and, we performed family-level RFX BMS as implemented in the VBA Toolbox. Family-level inference serves to reduce uncertainty about aspects of model structure other than the characteristic of interest (Penny et al., 2010). In the present study, we had no specific hypothesis about which of our candidate models explained the data best. Instead, we aimed to show that informing our response time models using quantities from the perceptual model (informed RT family) resulted in a better explanation of the data than using different variants of response time models that explain response times independent of the perceptual model (uninformed RT family). RFX BMS represents a hierarchical approach to model selection, treating the model as a random variable among subjects, allowing for inference on posterior family and model probabilities. RFX BMS accounts for group heterogeneity (different participants may be using different winning models/families) and provides robustness against outliers as opposed to fixed-effects (FFX) procedures (Stephan et al., 2009).

For the family-level RFX BMS, we specified a uniform prior at the family level to avoid biasing our inference (Penny et al., 2010). We computed the posterior family probabilities which correspond to the posterior belief that family *k* generated the data. Additionally, we computed family XPs which correspond to the belief that family *k* is more likely than any other of the *K* families, given the data from all participants.

**Aim 2**. A quantitative assessment of whether and how parameters characterising subject-specific learning behaviour were associated with an individual’s measured response times. To this end, we first needed to identify a single winning model – i.e. the model that best describes the measured data – by conducting a second RFX BMS analysis on the entire model space, this time at the level of individual models.

Once this model had been identified and in case it belonged to the family of informed RT models, we analysed the influence of the individual regressors of its response time GLM in a second step by examining the posterior parameter estimates of the winning model. To quantify the importance of each free parameter, we tested its significance against the *initial prior* mean (i.e. 0) using one-sample *t*-tests at a significance threshold of *p* < 0.05, Bonferroni-corrected for the number of performed tests (i.e. free parameters).

### Model evaluation

After we had fit our models to empirical data and performed a set of statistical tests to answer our research questions, it still remained to be evaluated whether our model(s) actually provided a good explanation of the data, i.e. an assessment of model quality in absolute terms, using posterior predictive checks (van de Schoot et al., 2021). This approach is distinct from the purpose of (Bayesian) model comparison which assesses the quality of candidate models in relative terms.

Specifically, we performed qualitative posterior predictive checks (van de Schoot et al., 2021) by simulating new data conditional on the obtained subject-specific posteriors. For this, we used the Hessian of the negative log joint at the MAP estimate obtained during the optimisation to determine the posterior covariance matrix. This allowed us to construct an approximate multi-dimensional posterior for each model and participant. We then sampled parameter values (*N*_*ppc*_ = 100) from the calculated approximate posterior densities of each participant and model and used the sampled parameter values to generate synthetic response data (binary and continuous). Subsequently, the generated data were compared to the empirical data at an individual subject level. In order to determine the quality of the posterior predictions of our model, we pre-specified the following criteria in our analysis plan which we deemed critical characteristics of the collected data set: Calculated adjusted correctness of binary responses, and the plausibility of predicted log RT trajectories assessed in a qualitative fashion.

## Results

### Model-agnostic results

Here, we report descriptive statistics of behavioural data from *N*_*main*_= 59 participants in our main sample. In Figure 3, mean and standard deviation of trial-wise binary responses and log RTs across participants are visualised in red. Figure 4A displays a trial-wise summary of percentage of incorrect responses over all participants. A histogram of log-transformed RTs is presented in Figure 4B. Behavioural data of individual participants can be found in the Supplementary Material (S1). A one-way ANOVA of log-transformed RTs did not reveal a significant main effect of factor phase (*p* = 0.15).

**Figure 4.**
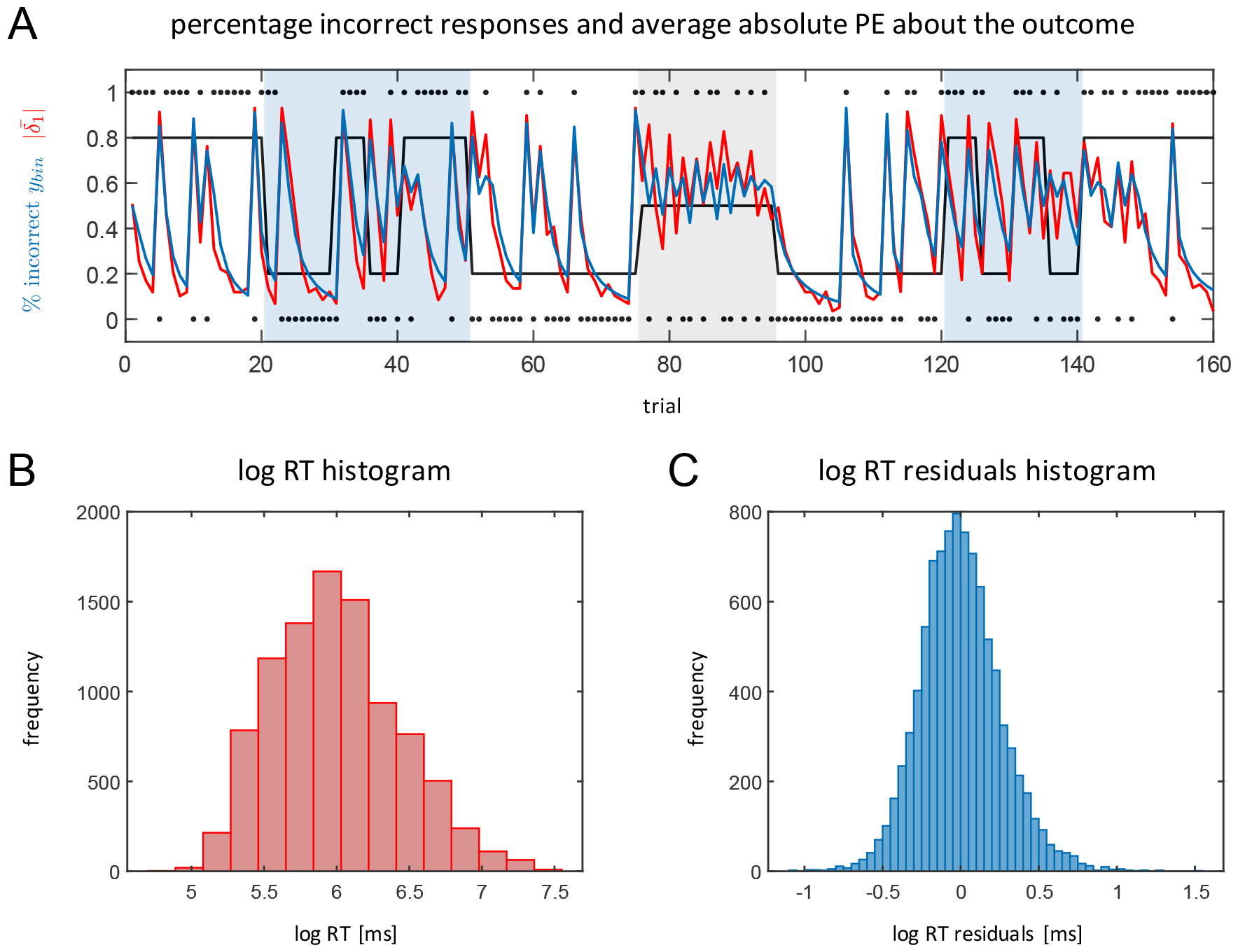
Binary responses and continuous log-transformed response times. In **A**, the red line represents a trial-wise summary of percentage of incorrect responses (inverted for true probabilities 0.2 indicated by the black line) over all participants (*N*_*main*_= 59) and the average absolute prediction error about the trial outcome 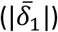 of M1 in red. Black dots represent the reward of fractal A on each trial (1=reward, 0=no reward) and the black line shows the underlying probability structure of the task. **B** shows the histogram of log-transformed RTs over all participants in ms. The histogram of residuals of log RT model fits obtained by M1 are visualised in **C**.

### Computational Modelling

#### Prior specification

A detailed treatment of the *initial priors* including their sufficient statistics, prior predictive distributions, and a grid search for different parameter values of the eHGF is presented in the analysis plan (Table 1, Figures 3 and A1). Here, we focus on the *empirical priors* which are used for the analysis of the main data set.

In Figure 5A *empirical prior* densities for all free parameters of M1 are shown alongside the MAP estimates from model inversion on the pilot data set (*N*_*pilot*_ = 20) as well as the *initial prior* densities. (Note that these reported *empirical priors* differ slightly from the specifications in the analysis plan, which is due to the fact that we are using a later version of TAPAS to run the analysis. However, qualitatively the *empirical priors* do not differ between the two versions.) It can be seen that the informativeness of the priors increased from the *initial* to the *empirical priors*. Again, it is worth emphasising that we avoided problems of circularity (i.e. informing the prior by the same data set in whose analysis the prior is applied), achieved by using an independent data set to estimate the sufficient statistics of the *empirical priors*.

**Figure 5.**
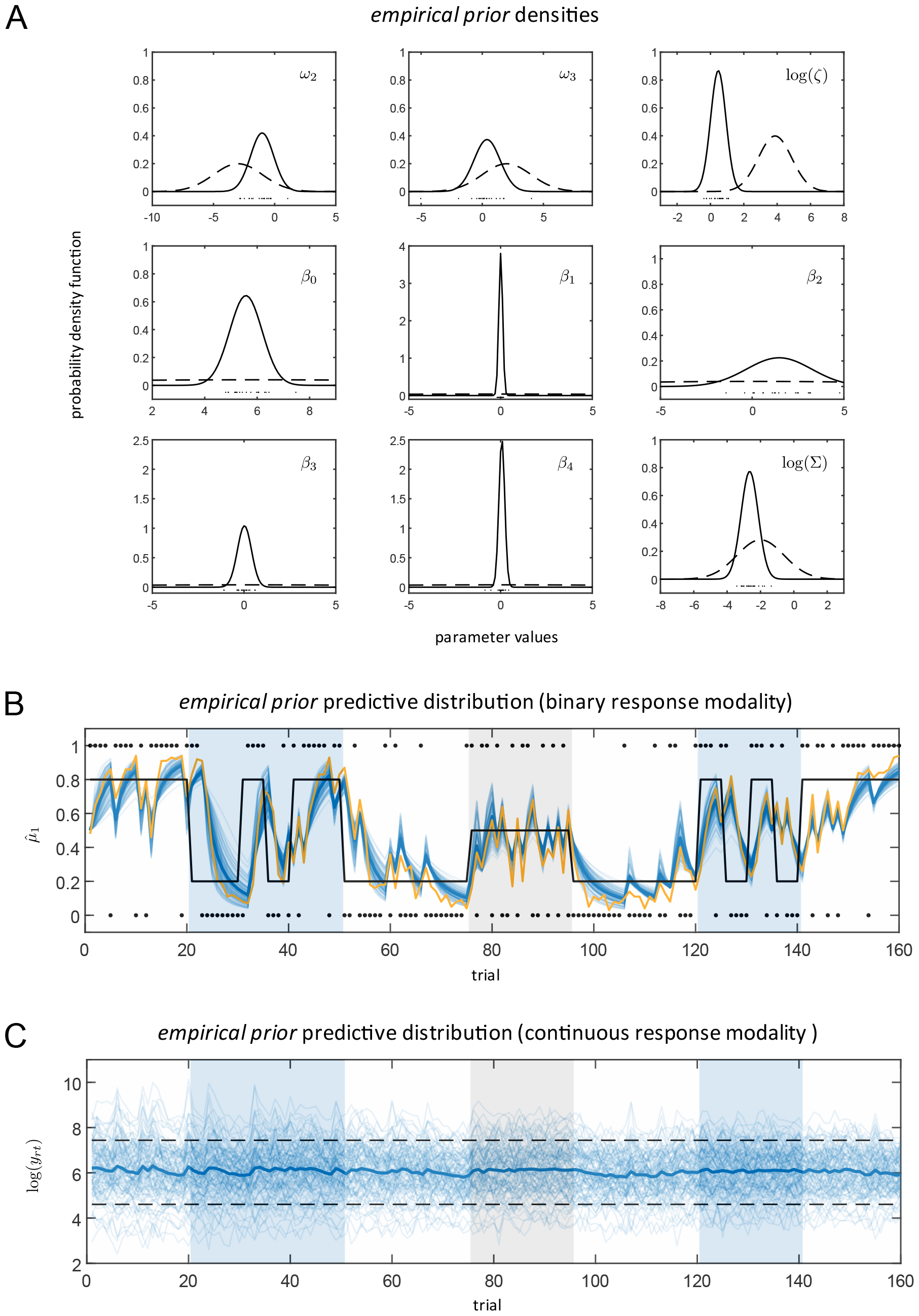
Prior configurations of M1. **A** shows the *empirical prior* densities for each free parameter of M1 (solid line) as estimated using MAP estimates (black dots) obtained from a separate pilot data set (*N*_*pilot*_ = 20) using the *initial priors* (dashed lines). A detailed description of M1 can be found in Table 1 and in the main text. Prior predictive distributions under the *empirical priors* of M1 are displayed for both response data modalities. In **B**, belief trajectories about the outcome 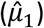 at the lowest level of the eHGF is displayed in blue with the thick blue line representing the resulting belief trajectory using the *empirical prior* mean parameter values and the yellow line representing the average simulated binary response (*N*_*sim*_ = 100). In **C**, simulated log RT data are shown in blue, the dashed black lines are the boundaries given by the length of the response window in each trial. The thick blue line represents the average simulated log RT trajectory (*N*_*sim*_ = 100).

Prior predictive checks using the *empirical priors* are visualized in Figures 5B and 5C. Figure 5B shows the distribution of predicted belief trajectories 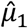 at the outcome level of the eHGF as well as the trial-wise frequency of simulated binary responses of M1 (*N*_*sim*_ = 100). Since we assume relatively little decision noise under the *empirical priors*, the frequency of binary responses is higher than the average predicted belief for trials where 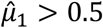 and vice-versa for trials where 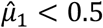. Figure 5C displays simulated log RT trajectories for M1 under the *empirical priors*. Visualisations of the *empirical prior* densities, the respective prior predictive distributions at all three levels of the perceptual model as well as simulated log RT data for all seven models in our model space are included in the Supplementary Material (S2).

Across models, our prior predictive checks using the *empirical priors* showed that – as expected by the construction of the model space – all our models produce similar behaviour at the level of the perceptual model and how they predict binary response data. By contrast, they differ considerably in the way they predict log RT data (see Supplementary Material Figures S2O-S2P). Moreover, all of them allow for a wide range of behaviour indicating that the elicited *empirical prior* densities are flexible enough to account for inter-individual differences between participants. This serves as qualitative sanity check of our prior elicitation procedure.

#### Validation of computation

Results from the family-level recovery analysis (Figure 6A) show good recoverability of true family frequencies for the 5’000 synthetic data sets (*N*_*famsim*_ = 60 simulated subjects per data set). Reassuringly, both Ef as well as XP values show little to no bias towards either of the two model families.

**Figure 6.**
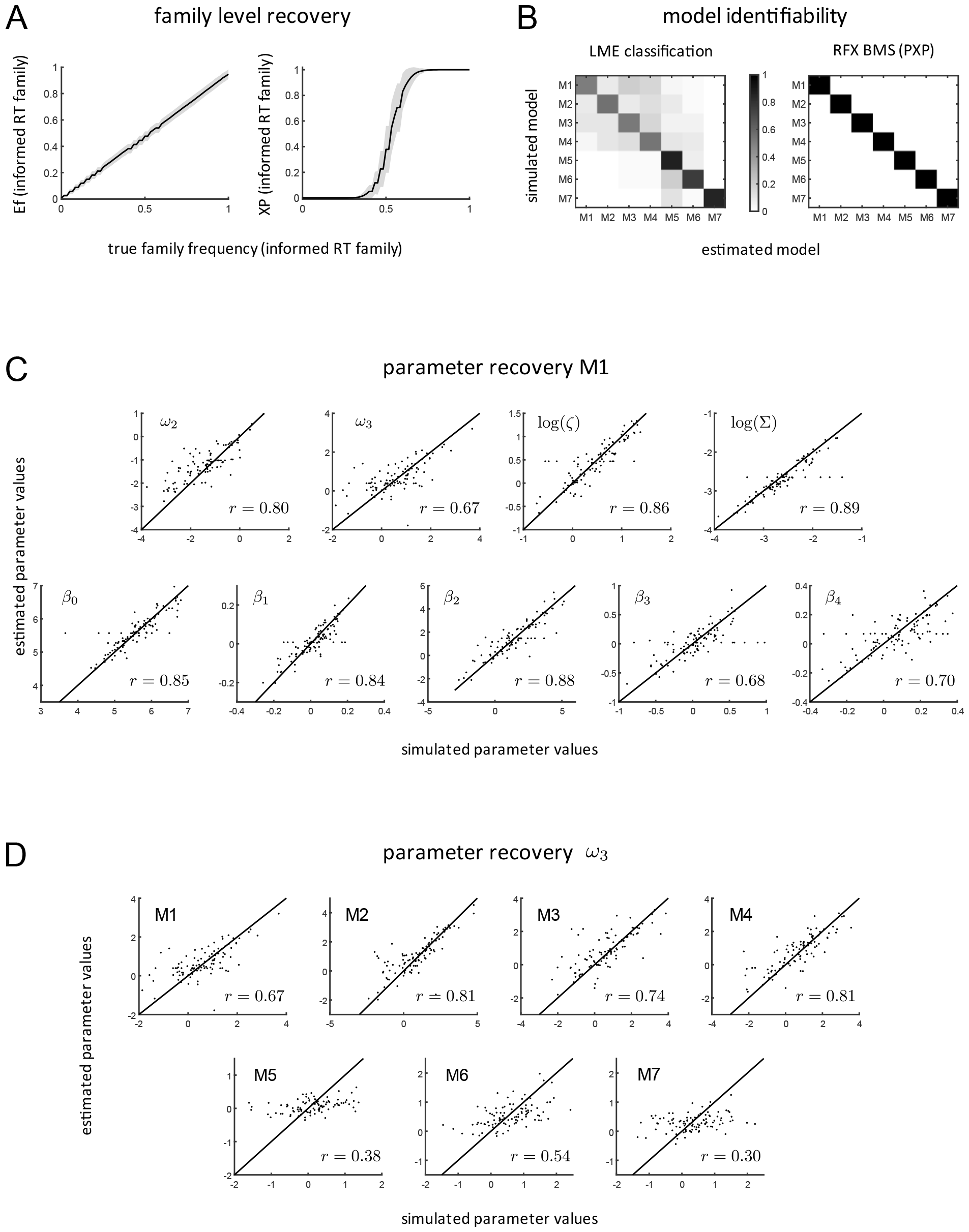
Validation of Computation. **A** shows results from family-level recovery analysis comparing Ef (left) and XP (right) values with true family frequencies. **B** depicts results from individual model level recovery analysis. 7x7 confusion matrices for LME winner frequencies (left) and PXP scores (right) are shown with data generating models on the y-axis and recovered models on the x-axis. **C** visualizes simulated vs. estimated values of all free parameters of M1 (parameter recovery). Correlation between simulated and estimated parameter values is indicated using Pearson correlation coefficients *r*. In **D**, recoverability of the *ω*_3_ parameter of the 3-level eHGF for binary inputs is visualised for all seven models in the model space including Pearson correlation coefficients *r*. Models from the informed RT family (M1-M4) show consistently better *ω*_3_ recovery compared to models of the uninformed RT family (M5-M7).

Figure 6B shows the 7x7 confusion matrices resulting from Individual-level model recovery analysis. All models could be identified well above chance level both when evaluating approximate LME winner frequencies as well as PXP values resulting from RFX BMS on the synthetic data set (*N*_*sim*_ = 100). For the LME winner classification analysis, the balanced accuracy score is 0.66 which is clearly above 0.19 (the upper bound of the 90%-CI when assuming chance across all 7 models).

Parameter recovery analysis of M1 is visualised in Figure 6C. All of the nine free parameters of M1 (two perceptual and seven response model parameters) show good recoverability. All Pearson correlations between true and recovered parameter values are highly significant (*p* < 0.001). Recoverability of the meta-volatility parameter (*ω*_3_) of the 3-level eHGF for binary inputs is the least robust among all free parameters of M1 (*r* = 0.67). Detailed results from parameter recovery analysis of all models are listed in the Supplementary Material (S3). Most of the model parameters are well recoverable in all of the seven models. Notably, in all models, the meta-volatility parameter (*ω*_3_) of the perceptual model is more challenging to recover than other parameters. However, Figure 6D shows that this depends on whether information about RTs is considered by the response model or not: models of the informed RT family (M1-M4) show much better *ω*_3_ recoverability (with correlation coefficients in the range of 0.67-0.81) compared to models of the uninformed RT family (M5-M7), where correlation coefficients are in the range of 0.3-0.54. This shows a clear benefit in terms of practical identifiability brought by the use of response models that combine different data modalities in a way that incorporates states from the perceptual model in both parts of the response model.

#### Model comparison

**Aim 1**. Figure 7A shows the results of the family level RFX BMS on the main data set. The family of informed RT models is identified as the winning family (*XP* = 1, *Ef* = 0.79). In other words, we find clear evidence supporting the hypothesis that the family of informed RT models explains the collected data better (trading off accuracy and complexity) compared to models from the uninformed RT family. This demonstrates the practical utility of the family of informed RT models.

**Figure 7.**
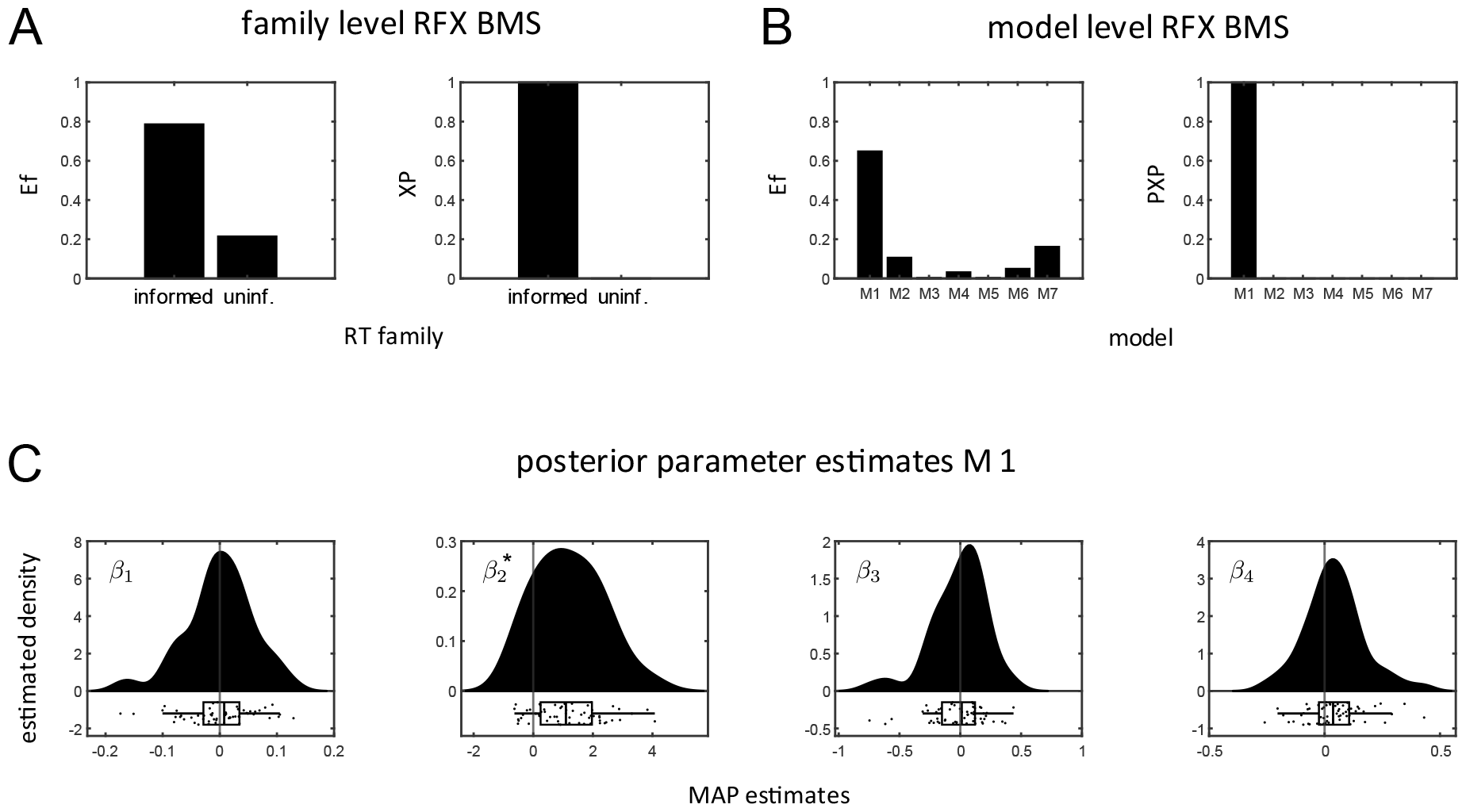
Hypothesis testing. **A** shows the results of the family level RFX BMS (Efs on the left, XPs on the right) with the informed RT family clearly outperforming the uninformed RT model family. **B** displays Efs (left) and PXPs (right) resulting from individual model level RFX BMS. M1 can be identified as the clear winning model. **C** shows raincloud plots of the MAP estimates of the M1 GLM regressors (generated using the RainCloudPlots library). Fine black vertical lines indicate the *initial prior* mean values (i.e. 0) and a black star indicates significantly different prior and posterior means of *β*_2_ which scales the influence of 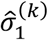 on log RTs (one-sample *t*-test, *p* < 0.001).

**Aim 2**. Figure 7B visualizes the results of the individual model level RFX BMS on the main data set. Here, M1 is identified as the winning model (*PXP* = 1, *Ef* = 0.65), which consists of a 3-level eHGF for binary inputs combined with the unit-square sigmoid model as binary part and the Lawson-inspired log RT GLM as continuous part of the response model. Figure 7C displays MAP estimates of the M1 log RT GLM regressors. Post-hoc one-sample *t*-tests on posterior means of the log RT GLM parameters of M1 reveal a highly significant (*p* < 0.001) difference between *initial prior* mean (i.e. 0) and posterior mean values of the parameter *β*_2_, which scales the influence of the informational uncertainty at the level of the outcome (Bernoulli variance 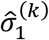) from the perceptual model on the log RTs. None of the other GLM regressors shows a significant difference between prior and posterior mean values (Bonferroni-corrected).

#### Model evaluation

Figure 3 shows that the average binary predictions as well as the average predicted log RTs by M1 over all participants of the main data set capture the empirical data quite well. In Figure 4A, the trial-wise percentage of participants giving an incorrect response (inverted for true reward probabilities of 0.2 to improve visual comparability) is compared to the averaged absolute prediction error at the outcome level 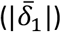 of M1. This qualitative comparison serves as an example of how the low-level prediction error computed using the 3-level eHGF for binary inputs resembles participants’ performance during the SPIRL task. Figure 4C displays the residuals of log RT model fits obtained by M1 which are approximately normally distributed. This is in accordance with the assumption of Gaussian noise in the log RT GLMs.

Examples of the best, average and worst fits of M1 to single-participant data in terms of the log likelihood are shown in the Supplementary Material (S1). Overall, belief trajectories at the outcome level of the eHGF seem to align well with the participants’ binary predictions. Regarding the RT fits, one can clearly see that individual log-transformed RT data is very noisy. However, a look at the averaged log RT data across participants and the averaged log RT model fits clearly shows that M1 is able to pick up the overarching structure in the log RT trajectories (Figure 3). Importantly, models from the informed RT family (M1-M4) and to some degree also M7 show a close correspondence between the average of the predicted and the average of the actual RT trajectories (see Supplementary Material S4).

Results from posterior predictive checks of M1 are shown in Figure 8 and in the Supplementary Material (S5). These allow for a qualitative assessment of the obtained single-subject posteriors. Regarding the binary response data, we can clearly see that the adjusted correctness of empirical response data is within the ranges of values covered by the individual subjects’ posterior predictive densities for most of the participants (Figure 8A and Supplementary Figure S5A). Concerning the response time data, we can observe that predicted signal using posterior mean parameter values captures higher level fluctuations in individual empirical log-transformed RT trajectories (Figure 8B and Supplementary Figure S5B). Similar to our observations regarding the individual-level RT model fits, sampled log RT trajectories are very noisy. However, the range of simulated log RT trajectories from the obtained posteriors aligns well with the individual empirical log RT trajectories. Moreover, the histograms of simulated log RT data from single-subject posteriors show a higher dispersion for participants with worse model fits as measured by the log likelihood (Figure 8B).

**Figure 8.**
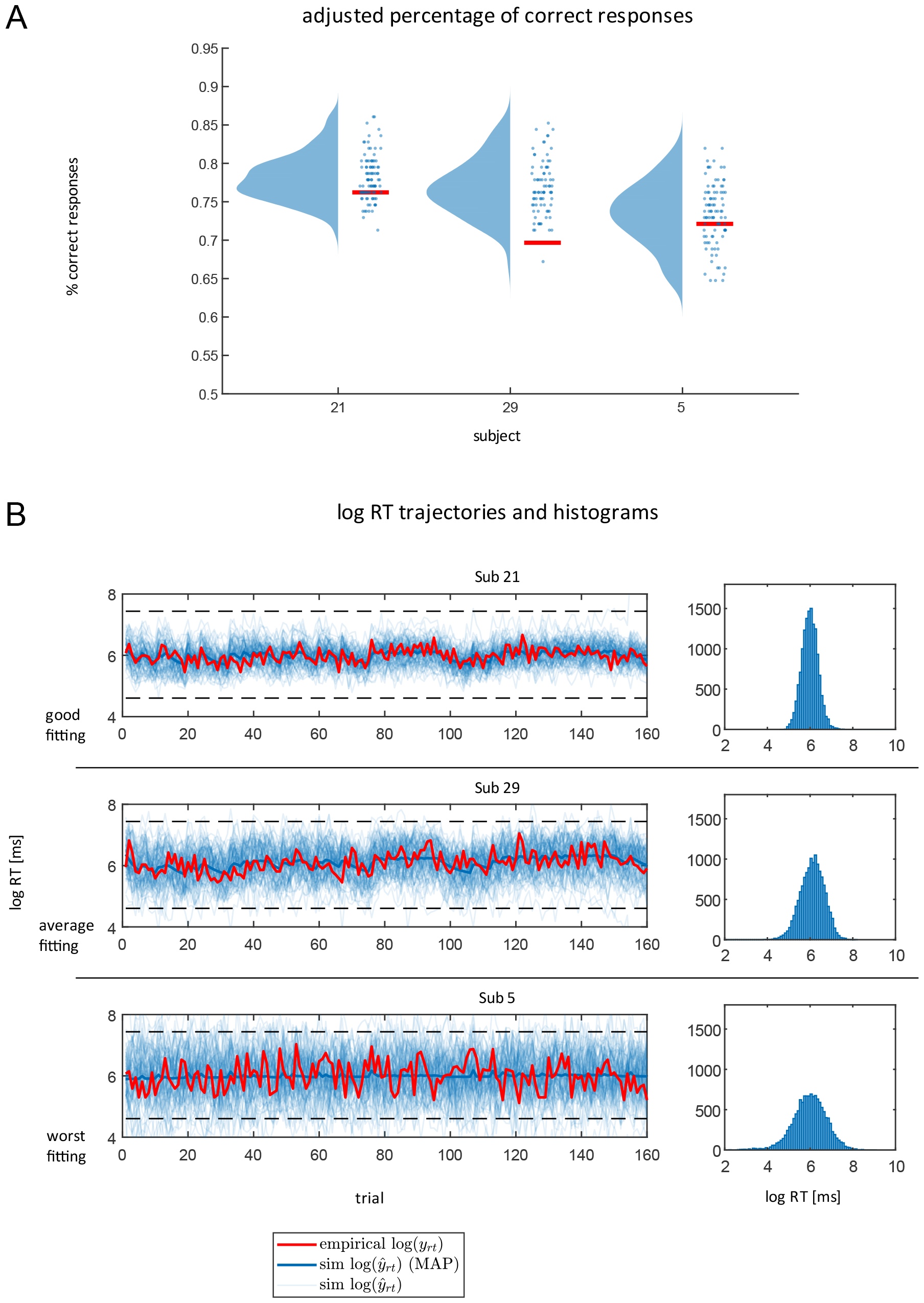
Posterior predictive checks for M1. Data from three participants of the main data set are shown. Participants were chosen according to goodness of model fit of M1, i.e. participant 21 with a high log likelihood value, participant 29 showing average goodness of fit and participant 5 showing the worst fit. **A** displays adjusted correctness of binary responses for these participants in red. Blue circles are the simulated adjusted correctness values resulting from sampled parameter values of the subject-specific posteriors of M1. The blue probability densities are the estimated posterior predictive densities based on the samples drawn from the posteriors (*N*_*ppc*_ = 100) using kernel density estimation as implemented in the RainCloudPlots library. In **B**, we show empirical log RT trajectories of the three participants in red. Fine blue lines are simulated log RT trajectories resulting from sampled parameter values of the subject-specific posteriors and the thick blue line represents the predicted log RT when using the MAP estimates of M1 for each participant to generate synthetic RT data. The histograms on the right visualise the distribution of synthetic log RT data generated by simulating from the subject-specific posteriors.

## Discussion

The present study provides a first application of response models combining two different data modalities in the framework of the HGF. More specifically, we developed a novel set of response models simultaneously fitting binary responses and continuous response time data during inference. Additionally, we developed and implemented an associative learning task, the SPIRL task, in which fast responses were incentivised, allowing us to model reaction times and binary responses jointly. We performed extensive simulation analyses and applied our set of combined response models to behavioural data from the SPIRL task. Our computational approach highlighted the utility of Bayesian workflow, increasing transparency and interpretability of reported results. We demonstrated the advantage of combining different response data modalities in a single model for the robustness of inference. Finally, the analysis of the data from the SPIRL task resulted in a clearly superior model, providing the best explanation of the data. Inspection of individual parameter estimates of this model revealed a significant linear relationship between log RT data and informational uncertainty at the level of the outcome 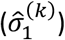.

### Combining different response data modalities for robust inference in the HGF

The first motivation for this paper was the development and application of a set of response models that combined multiple data modalities. In this way, we hoped to address issues related to parameter recovery and model identifiability in the context of the HGF, especially when dealing with applications to binary response data (Harrison et al., 2021; Iglesias et al., 2021). We hypothesised that the combination of different data modalities should increase identifiability, both at the level of parameters and models. Recoverability of parameters should benefit further from incorporation of perceptual model quantities in the response models. Indeed, our simulations demonstrated good to excellent recoverability at all different levels (model parameters, individual models, and model families). We also showed that recoverability of perceptual model parameters was superior in the family of informed RT models, which use quantities from the perceptual model as part of the log RT GLM, as opposed to the family of uninformed RT models.

Comparing the performance of the two model families against empirical data and using family level RFX BMS, we demonstrated that the data from the SPIRL task were better explained by the family of informed RT compared to the family of uninformed RT models. Moreover, RFX BMS on the individual model level revealed M1 as the clearly winning model, providing the best explanation for the empirical data set.

Our analysis of the regression weights of the log RT GLM of M1 revealed a significant influence of the parameter *β*_2_ which scales the contribution of the outcome uncertainty 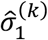 in the eHGF. Here, we refer to outcome uncertainty as the informational uncertainty at the level of the outcome, which describes the irreducible uncertainty associated with any type of probabilistic prediction. This result is consistent with the findings of Lawson et al. (2017) who applied a very similar combination of an HGF perceptual model and a GLM as response model, mapping states of the perceptual model to log RT data in an associative learning task. In addition, other work that did not use the HGF framework also found evidence for a relationship between reaction times and the uncertainty of responses (Bonnet & Ars, 2008).

Harnessing the information of an additional data modality during statistical inference is not a novel idea. First examples of similar approaches exist in the context of sequential sampling models such as drift-diffusion models (DDMs) (Kraemer et al., 2021; Pedersen et al., 2017; Shahar et al., 2019) which have been combined with RL models (Ballard & McClure, 2019; Clithero, 2018; Loeys et al., 2011; McDougle & Collins, 2021; Miletić et al., 2021. However, in the literature, models combining binary choices and continuous response times are far less common than simple binary observation models. Moreover, our study focuses on a specific generative modelling framework, the HGF, for which the present study, to the best of our knowledge, represents the first application of a response model combining different response data modalities to empirical data in combination with the HGF.

There are some limitations to our modelling approach. First of all, for the specification of our model space we assumed independence between the two response data modalities (binary responses and response times) conditional on the parameters of the perceptual model. For future applications, one may consider finessing the current formulation. Second, MAP inference is a rather simplistic technique for Bayesian inference where the posterior uncertainty needs to be approximated post-hoc. Alternatives would be VB or MCMC methods which directly include an estimate of the posterior uncertainty. Moreover, the chosen inference method is based on a variant of gradient descent, which might not be optimal for dealing with multimodal posteriors. We tried to combat this shortcoming by adopting a multistart approach to prevent the optimisation from getting stuck at local extrema. The results from parameter recovery analysis suggested that the chosen inference scheme is appropriate for the application at hand.

The presented combined response models for the HGF framework have potential for applications in a variety of tasks and domains. In principle, our modelling approach can be applied to any two different data modalities of interest, e.g. behavioural, physiological, neurophysiological data, etc. However, it is important that the data modalities of interest contain relevant information that can be picked up by the model. This can be demonstrated e.g. by comparing different models with a null model in a candidate application, similar to the comparison of different model families in this study.

We hope that the use of combined response models can be particularly useful in TN/CP. Typical constraints for the design of clinically applicable paradigms include limited number of trials and complexity constraints for tasks, which limit the number of data points and ultimately constrain inference. Added information from a second data modality may compensate for these limitations and thus help improve the robustness of results.

### Bayesian workflow for generative modelling in TN/CP

The second motivation for this paper was to highlight the key ingredients of Bayesian workflow and illustrate its application. In TN/CP, generative models not only represent central tools for inference on disease-relevant cognitive and neurophysiological mechanisms (Stephan & Mathys, 2014) but also frequently serve to provide low-dimensional and mechanistically interpretable features for machine learning (generative embedding; Stephan et al., 2017). The robustness of results from generative modelling is therefore of major importance in TN/CP and can benefit from incorporating general (field-unspecific) methodological and conceptual developments concerning BDA.

In BDA, the choice of priors plays an important role for identifiability, reliability and predictive validity of model-fitting results (Gershman, 2016). Yet, the importance of priors is easily overlooked and little attention is usually devoted to systematic description and analysis thereof. Similarly, validation of the chosen inference technique (including the effects of approximations and the choice of optimisation algorithms) as well as model evaluation are often neglected. This is especially critical in TN/CP, since the potential success of computational modelling endeavours for clinical applications is inextricably tied to the robustness of inference. Hence, transparency with regard to the hyperparameters of an analysis pipeline is important and requires a detailed description of individual analysis steps.

Importantly, we did not invent the Bayesian workflow presented here; instead, it was derived from previous proposals (e.g. Betancourt, 2020; Gelman et al., 2020; Schad et al., 2020; van de Schoot et al., 2021) and augmented by additional components, e.g. Bayesian model selection at the family level (Penny et al., 2010). These steps are summarised visually in Figure 2 and include the specification of an initial model space, prior elicitation and prior predictive checking, the choice of a Bayesian inference algorithm and concurrent validation of computation, model comparison procedures, and model evaluation.

Using the well-known Bayesian workflow by Gelman and colleagues (2020) for comparison, our approach introduces several extensions. First, our procedure for elicitation of prior distributions involves an independent data set, allowing us to obtain a set of data-informed priors representative of actual human behaviour while at the same time avoiding problems of double-dipping. Second, the analysis was pre-specified in its entirety, (the only change concerned using a more current version of TAPAS, a deviation which we explicitly mention above). This pre-specification is somewhat in contradiction to the iterative procedure proposed by Gelman and colleagues, but we consider this a strength of our approach. Given the many degrees of freedom and the numerous cognitive biases that scientists may inadvertently be affected by, pre-registration is an important and effective protection for researchers against fooling themselves (Nosek et al., 2018). Our point is not that iterative model building should not be part of Bayesian workflow; however, we believe that it is important to combine it with ‘guard railings’ (such as a preregistered analysis plan). Additionally, whenever possible, independent data sets should be used, both for specifying priors and for evaluating the generalisability of the obtained results. We appreciate that this may not always be possible, for example, in situations where data sets result from rare, or even unique, events.

We are aware that the proposed Bayesian workflow is not perfectly generalizable to every application of generative models in TN/CP. It should rather be seen as a blueprint that can be adapted and extended to specific cases of generative modelling. Furthermore, there are important elements of Bayesian workflow which we did not implement and discuss here in detail. For example, there are principled (but computationally expensive) approaches to validating Bayesian inference algorithms such as simulation-based calibration (SBC) (Talts et al., 2018). Also, when using sampling-based approximations to the posterior, additional diagnostics (e.g. concerning convergence) are required. Previous papers on BDA provide detailed discussions of these (and other) topics (Betancourt, 2020; Gelman et al., 2013, 2020; Schad et al., 2020; van de Schoot et al., 2021).

In summary, this paper provided an illustrative application of Bayesian workflow in the context of an associative learning task that allowed for simultaneously modelling two behavioural readouts. We hope that this example will help pave the way towards standard adoption of Bayesian workflow and contributes to efforts of improving the transparency and robustness of results in TN/CP.

## Authors’ contributions

AJH, conceptualisation, data curation, formal analysis, investigation, project administration, methodology, software, visualisation, writing – original draft, writing – review and editing, publication of data and code; SI, conceptualisation, data curation, project administration, methodology, supervision, writing – review and editing; LK, SM, data curation, investigation, writing – review and editing; MMS, LR, CM, conceptualisation, methodology, writing – review and editing; JH, conceptualisation, methodology, code review, writing – review and editing, publication of data and code; OKH, SF, conceptualisation, methodology, supervision, writing – review and editing; KES, resources, conceptualisation, methodology, funding acquisition, supervision, writing – review and editing;

