## Supplementary Material for "Bayesian Workflow for Generative Modeling in Computational Psychiatry"

### S1. Examples of individual participants' behavioural data and fits of M1 (main data set)

Figures S1A-S1D show behavioural data and model fits (M1) of the four participants for which M1 fit the data best as measured by the log likelihood.

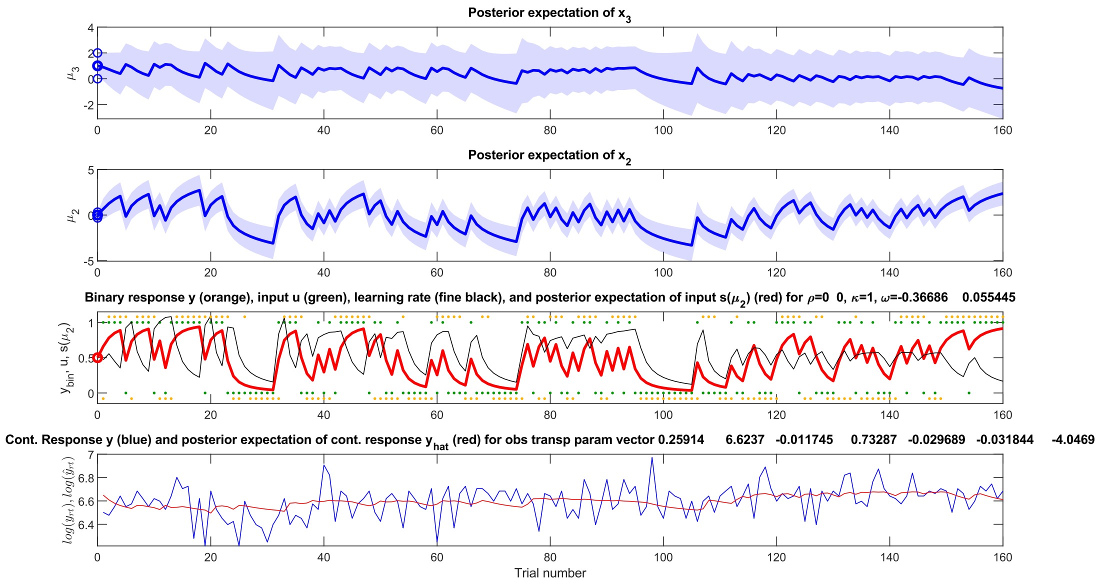


**Figure S1A** | Behavioural data and model fits (M1) of participant 45 of the main data set. The top panel shows the fitted mean belief $\mu_3$ about the log volatility of the environment in blue with the blue shaded area representing the uncertainty about $x_3$, i.e., the variance $\sigma_3$. The second panel shows mean $\mu_2$ and variance $\sigma_2$ of the belief about the cue-outcome contingency. The second lowest panel shows in red the mean belief about the outcome $\hat{\mu}_1$ given one of the two fractals, the actual outcomes for one of the two fractals as green dots (1 = reward, 0 = no reward) and which of the two fractals was selected as yellow dots. The fine black line represents an implied learning rate at the level of the outcome calculated as $\mathbb{1}_{\{\delta_1 \neq 0\}}\left(\frac{\Delta s(\mu_2)}{u-\Delta s(\mu_2)}\right)$ where $s$ is the sigmoid function. The lowest panel shows the empirical log RTs [ms] as blue line and the fitted log RT trajectory in red.

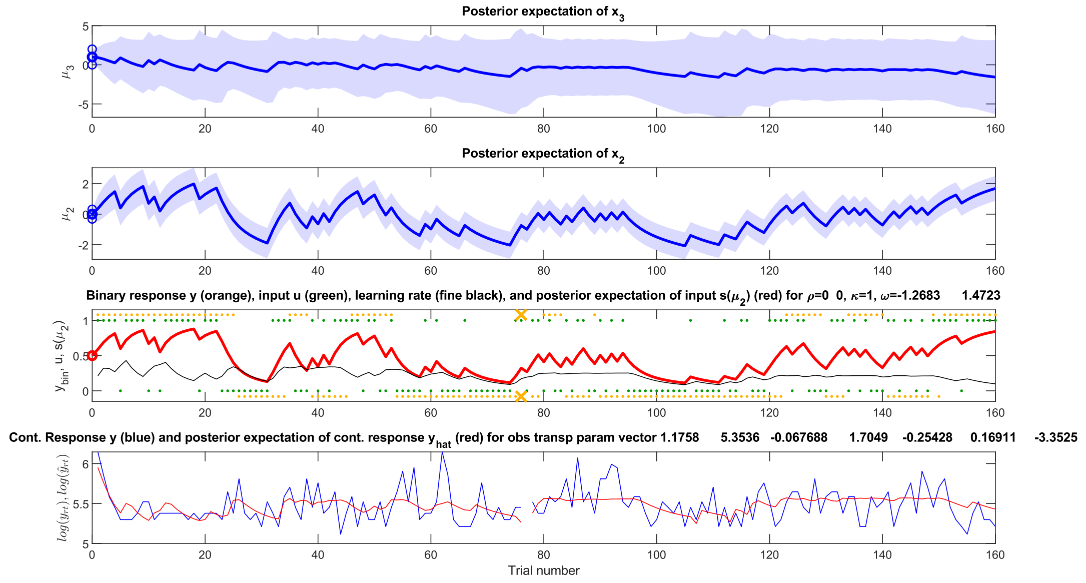


**Figure S1B** | Behavioural data and model fits (M1) of participant 3 of the main data set. The top panel shows the fitted mean belief $\mu_3$ about the log volatility of the environment in blue with the blue shaded area representing the uncertainty about $x_3$, i.e., the variance $\sigma_3$. The second panel shows mean $\mu_2$ and variance $\sigma_2$ of the belief about the cue-outcome contingency. The second lowest panel shows in red the mean belief about the outcome $\hat{\mu}_1$ given one of the two fractals, the actual outcomes for one of the two fractals as green dots (1 = reward, 0 = no reward) and which of the two fractals was selected as yellow dots. The fine black line represents an implied learning rate at the level of the outcome calculated as $\mathbb{1}_{\{\delta_1 \neq 0\}}\left(\frac{\Delta s(\mu_2)}{u - \Delta s(\mu_2)}\right)$ where $s$ is the sigmoid function. The lowest panel shows the empirical log RTs [ms] as blue line and the fitted log RT trajectory in red.

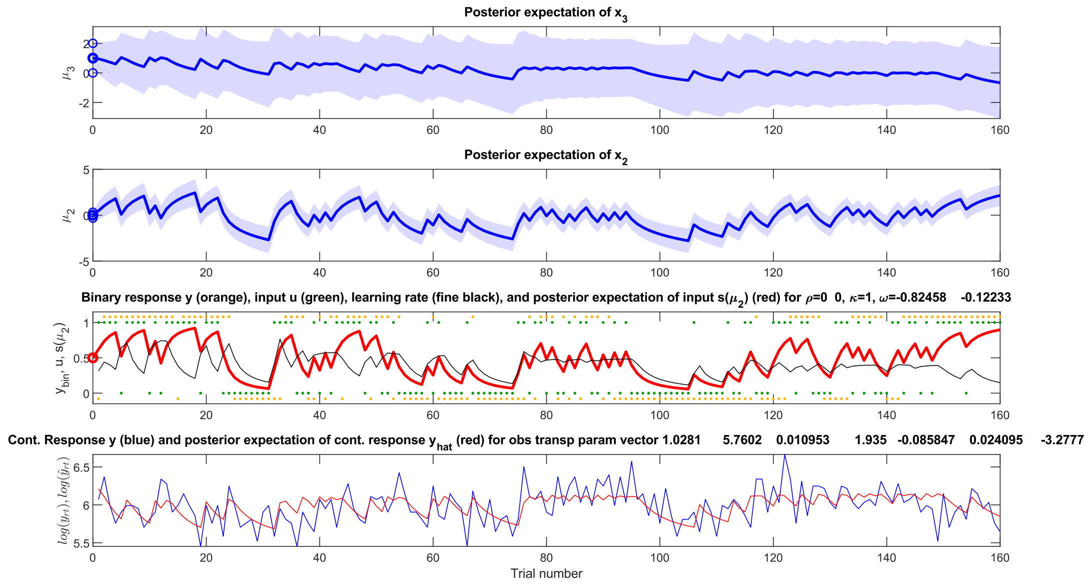


**Figure S1C** | Behavioural data and model fits (M1) of participant 21 of the main data set. The top panel shows the fitted mean belief $\mu_3$ about the log volatility of the environment in blue with the blue shaded area representing the uncertainty about $x_3$, i.e., the variance $\sigma_3$. The second panel shows mean $\mu_2$ and variance $\sigma_2$ of the belief about the cue-outcome contingency. The second lowest panel shows in red the mean belief about the outcome $\hat{\mu}_1$ given one of the two fractals, the actual outcomes for one of the two fractals as green dots (1 = reward, 0 = no reward) and which of the two fractals was selected as yellow dots. The fine black line represents an implied learning rate at the level of the outcome calculated as $\mathbb{1}_{\{\delta_1 \neq 0\}} \left(\frac{\Delta s(\mu_2)}{u - \Delta s(\mu_2)}\right)$ where $s$ is the sigmoid function. The lowest panel shows the empirical log RTs [ms] as blue line and the fitted log RT trajectory in red.

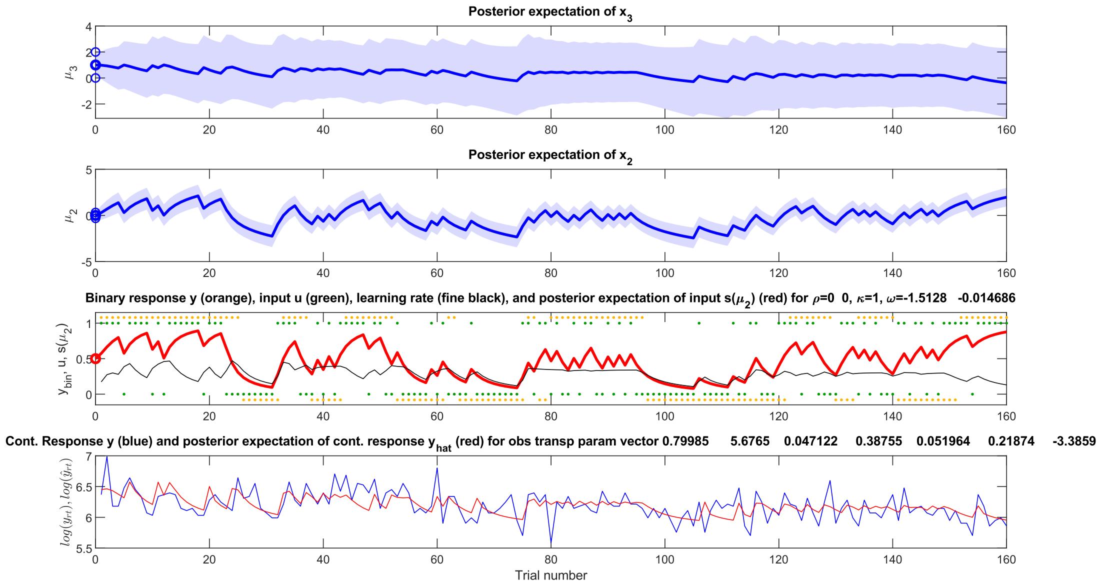


**Figure S1D** | Behavioural data and model fits (M1) of participant 33 of the main data set. The top panel shows the fitted mean belief $\mu_3$ about the log volatility of the environment in blue with the blue shaded area representing the uncertainty about $x_3$, i.e., the variance $\sigma_3$. The second panel shows mean $\mu_2$ and variance $\sigma_2$ of the belief about the cue-outcome contingency. The second lowest panel shows in red the mean belief about the outcome $\hat{\mu}_1$ given one of the two fractals, the actual outcomes for one of the two fractals as green dots (1 = reward, 0 = no reward) and which of the two fractals was selected as yellow dots. The fine black line represents an implied learning rate at the level of the outcome calculated as $\mathbb{1}_{\{\delta_1 \neq 0\}} \left(\frac{\Delta s(\mu_2)}{u - \Delta s(\mu_2)}\right)$ where $s$ is the sigmoid function. The lowest panel shows the empirical log RTs [ms] as blue line and the fitted log RT trajectory in red.

Figures S1E and S1F show behavioural data and model fits (M1) of two participants that show an average fit as measured by the log likelihood.

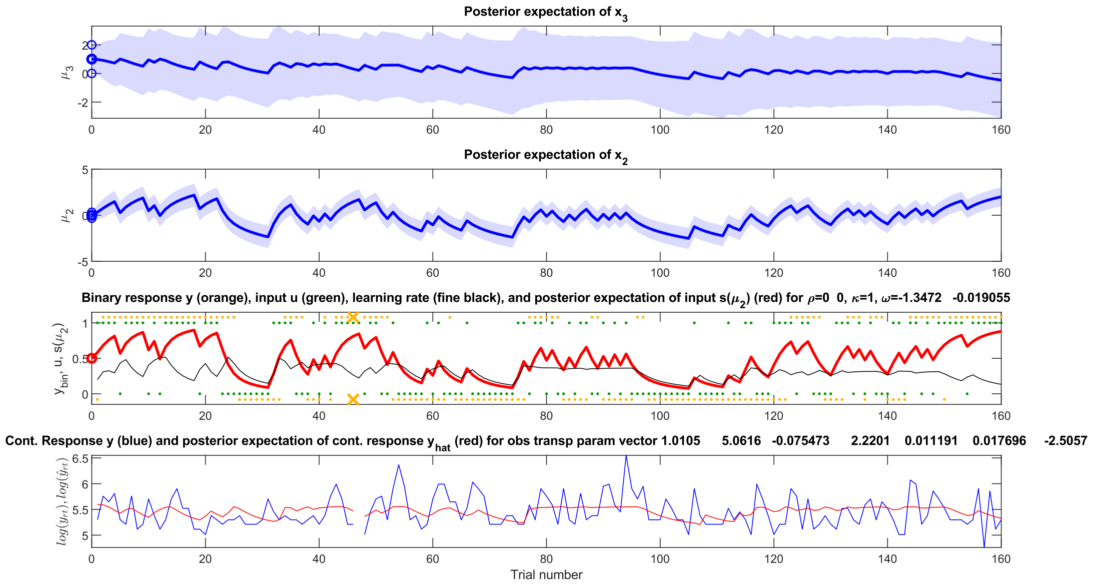


**Figure S1E** | Behavioural data and model fits (M1) of participant 28 of the main data set. The top panel shows the fitted mean belief $\mu_3$ about the log volatility of the environment in blue with the blue shaded area representing the uncertainty about $x_3$, i.e., the variance $\sigma_3$. The second panel shows mean $\mu_2$ and variance $\sigma_2$ of the belief about the cue-outcome contingency. The second lowest panel shows in red the mean belief about the outcome $\hat{\mu}_1$ given one of the two fractals, the actual outcomes for one of the two fractals as green dots (1 = reward, 0 = no reward) and which of the two fractals was selected as yellow dots. The fine black line represents an implied learning rate at the level of the outcome calculated as $\mathbb{1}_{\{\delta_1 \neq 0\}}\left(\frac{\Delta s(\mu_2)}{u-\Delta s(\mu_2)}\right)$ where $s$ is the sigmoid function. The lowest panel shows the empirical log RTs [ms] as blue line and the fitted log RT trajectory in red.

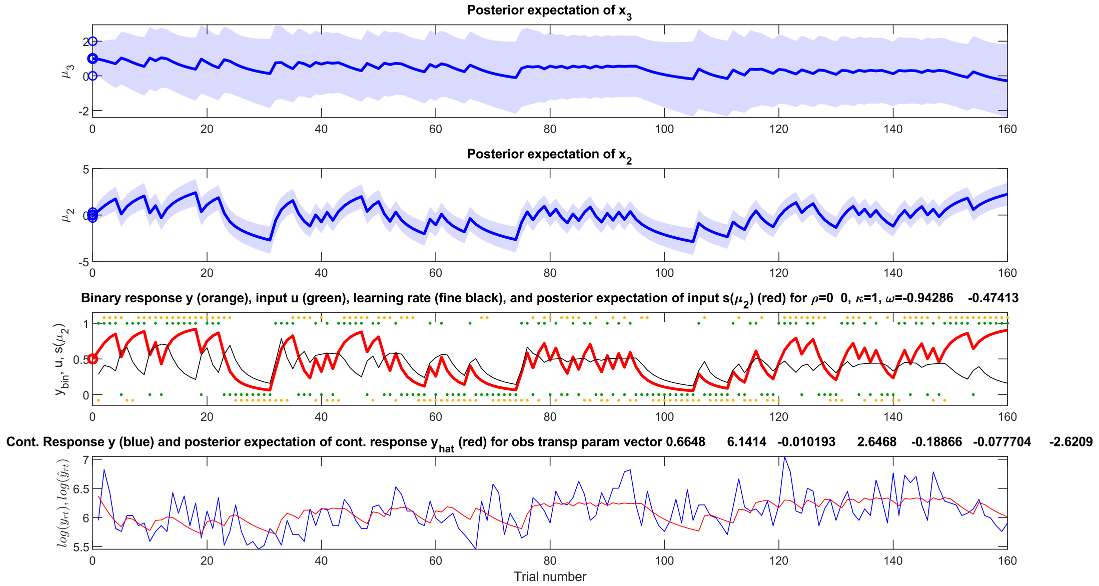


**Figure S1F** | Behavioural data and model fits (M1) of participant 29 of the main data set. The top panel shows the fitted mean belief $\mu_3$ about the log volatility of the environment in blue with the blue shaded area representing the uncertainty about $x_3$, i.e., the variance $\sigma_3$. The second panel shows mean $\mu_2$ and variance $\sigma_2$ of the belief about the cue-outcome contingency. The second lowest panel shows in red the mean belief about the outcome $\hat{\mu}_1$ given one of the two fractals, the actual outcomes for one of the two fractals as green dots (1 = reward, 0 = no reward) and which of the two fractals was selected as yellow dots. The fine black line represents an implied learning rate at the level of the outcome calculated as $\mathbb{1}_{\{\delta_1 \neq 0\}} \left(\frac{\Delta s(\mu_2)}{u - \Delta s(\mu_2)}\right)$ where $s$ is the sigmoid function. The lowest panel shows the empirical log RTs [ms] as blue line and the fitted log RT trajectory in red.

Figures S1G-S1J show behavioural data and model fits (M1) of the four participants for which M1 showed the worst fit of the data as measured by the log likelihood.

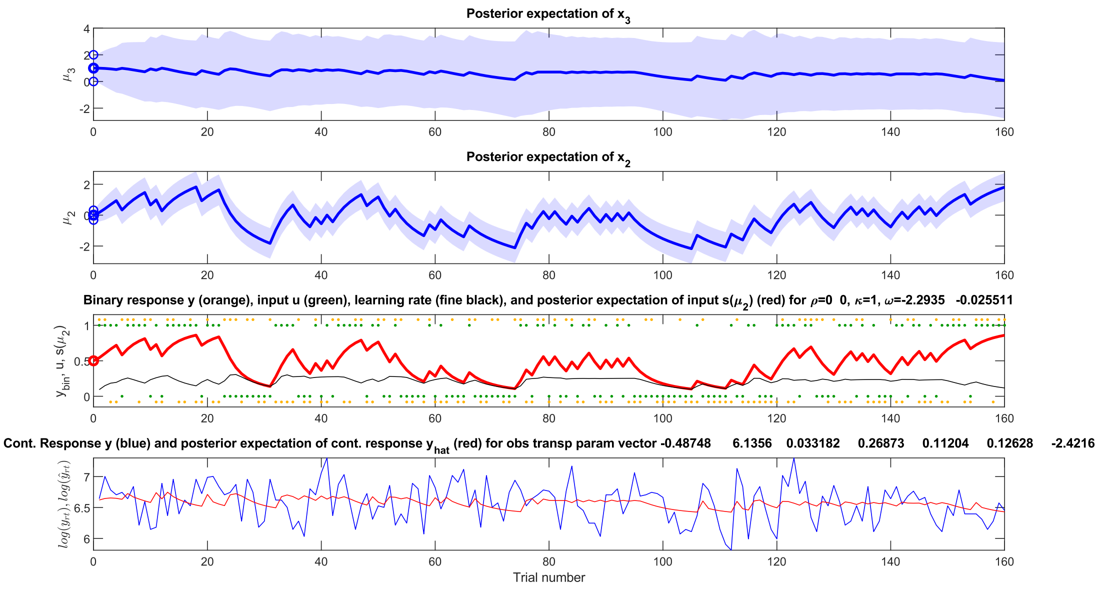


**Figure S1G** | Behavioural data and model fits (M1) of participant 6 of the main data set. The top panel shows the fitted mean belief $\mu_3$ about the log volatility of the environment in blue with the blue shaded area representing the uncertainty about $x_3$, i.e., the variance $\sigma_3$. The second panel shows mean $\mu_2$ and variance $\sigma_2$ of the belief about the cue-outcome contingency. The second lowest panel shows in red the mean belief about the outcome $\hat{\mu}_1$ given one of the two fractals, the actual outcomes for one of the two fractals as green dots (1 = reward, 0 = no reward) and which of the two fractals was selected as yellow dots. The fine black line represents an implied learning rate at the level of the outcome calculated as $\mathbb{1}_{\{\delta_1 \neq 0\}} \left(\frac{\Delta s(\mu_2)}{u - \Delta s(\mu_2)}\right)$ where $s$ is the sigmoid function. The lowest panel shows the empirical log RTs [ms] as blue line and the fitted log RT trajectory in red.

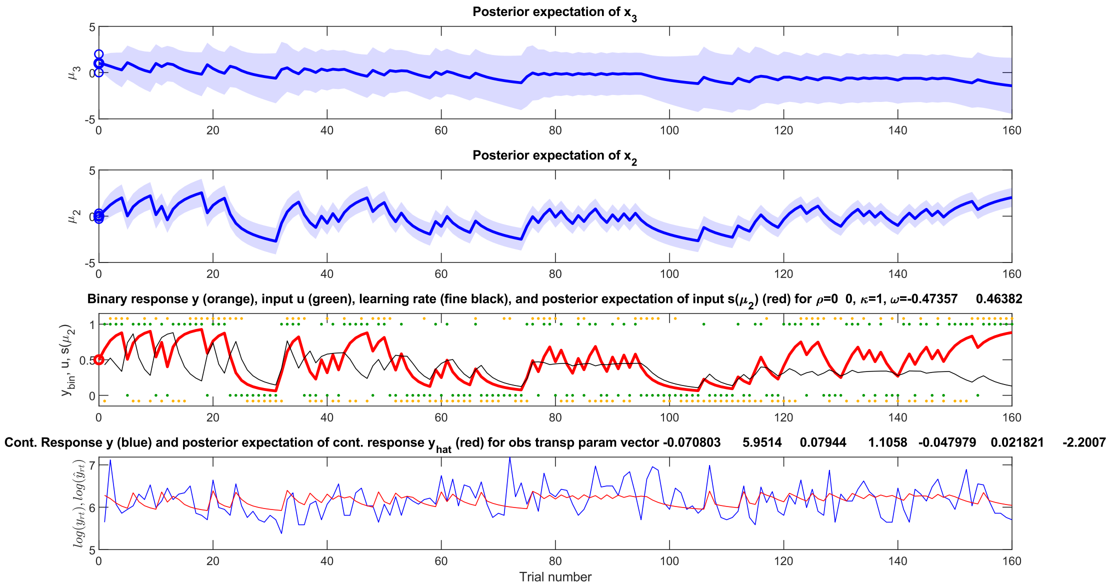


**Figure S1H** | Behavioural data and model fits (M1) of participant 18 of the main data set. The top panel shows the fitted mean belief $\mu_3$ about the log volatility of the environment in blue with the blue shaded area representing the uncertainty about $x_3$, i.e., the variance $\sigma_3$. The second panel shows mean $\mu_2$ and variance $\sigma_2$ of the belief about the cue-outcome contingency. The second lowest panel shows in red the mean belief about the outcome $\hat{\mu}_1$ given one of the two fractals, the actual outcomes for one of the two fractals as green dots (1 = reward, 0 = no reward) and which of the two fractals was selected as yellow dots. The fine black line represents an implied learning rate at the level of the outcome calculated as $\mathbb{1}_{\{\delta_1 \neq 0\}} \left( \frac{\Delta s(\mu_2)}{u - \Delta s(\mu_2)} \right)$ where $s$ is the sigmoid function. The lowest panel shows the empirical log RTs [ms] as blue line and the fitted log RT trajectory in red.

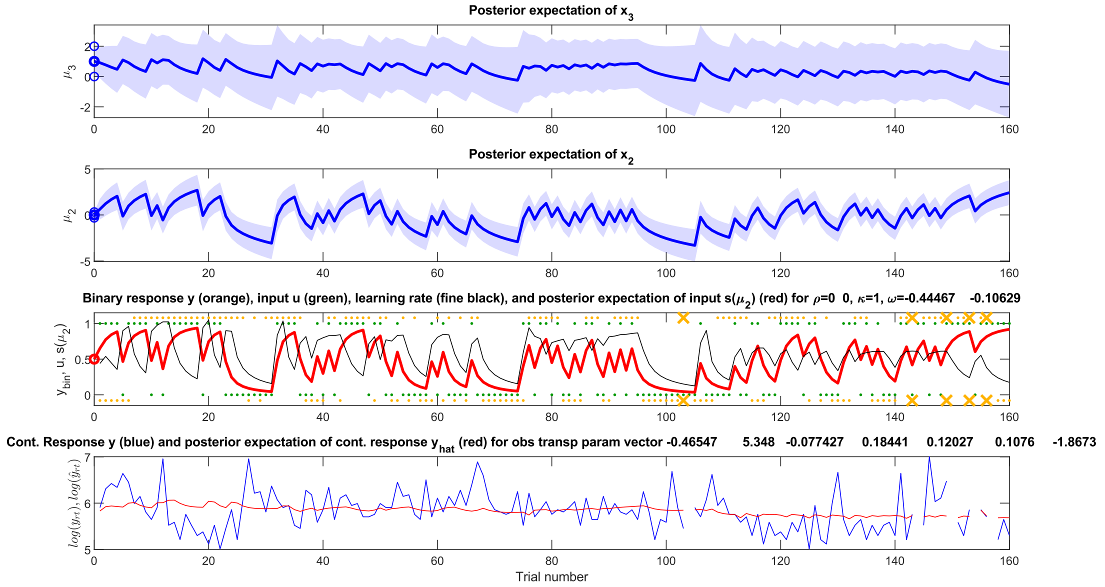


**Figure S1I** | Behavioural data and model fits (M1) of participant 15 of the main data set. The top panel shows the fitted mean belief $\mu_3$ about the log volatility of the environment in blue with the blue shaded area representing the uncertainty about $x_3$, i.e., the variance $\sigma_3$. The second panel shows mean $\mu_2$ and variance $\sigma_2$ of the belief about the cue-outcome contingency. The second lowest panel shows in red the mean belief about the outcome $\hat{\mu}_1$ given one of the two fractals, the actual outcomes for one of the two fractals as green dots (1 = reward, 0 = no reward) and which of the two fractals was selected as yellow dots. The fine black line represents an implied learning rate at the level of the outcome calculated as $\mathbb{1}_{\{\delta_1 \neq 0\}}\left(\frac{\Delta s(\mu_2)}{u-\Delta s(\mu_2)}\right)$ where $s$ is the sigmoid function. The lowest panel shows the empirical log RTs [ms] as blue line and the fitted log RT trajectory in red.

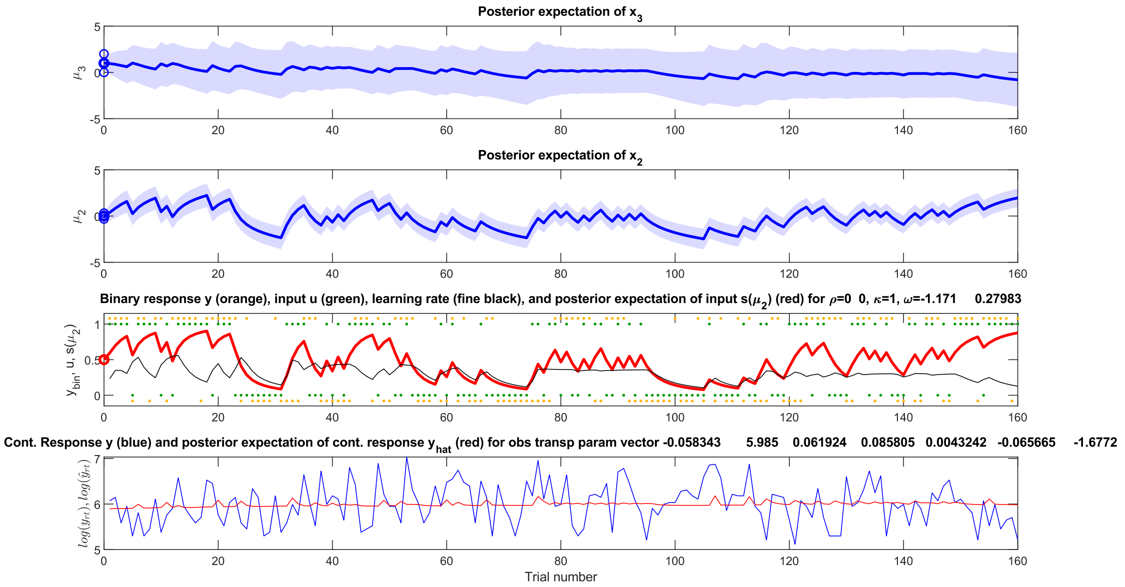


**Figure S1J** | Behavioural data and model fits (M1) of participant 5 of the main data set. The top panel shows the fitted mean belief $\mu_3$ about the log volatility of the environment in blue with the blue shaded area representing the uncertainty about $x_3$, i.e., the variance $\sigma_3$. The second panel shows mean $\mu_2$ and variance $\sigma_2$ of the belief about the cue-outcome contingency. The second lowest panel shows in red the mean belief about the outcome $\hat{\mu}_1$ given one of the two fractals, the actual outcomes for one of the two fractals as green dots (1 = reward, 0 = no reward) and which of the two fractals was selected as yellow dots. The fine black line represents an implied learning rate at the level of the outcome calculated as $\mathbb{1}_{\{\delta_1 \neq 0\}}\left(\frac{\Delta s(\mu_2)}{u-\Delta s(\mu_2)}\right)$ where $s$ is the sigmoid function. The lowest panel shows the empirical log RTs [ms] as blue line and the fitted log RT trajectory in red.

### S2. *Empirical priors*

#### S2a. *Empirical prior densities of M1-M7*

Both *initial* and *empirical prior* densities for all seven models are shown in Figures S2A-S2G.

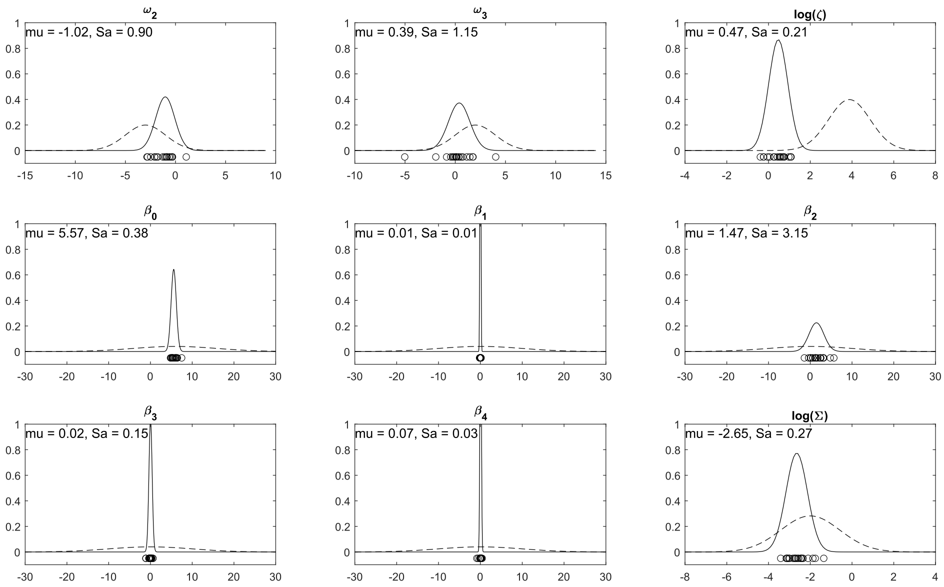


**Figure S2A** | *Empirical prior* densities of M1. For each free parameter of the model, the *initial prior* densities are shown as dashed line and the MAP estimates obtained from model inversion on the held-out data set ($N_{pilot}$) under the *initial priors* as black circles. The estimated *empirical priors* are shown as black line.

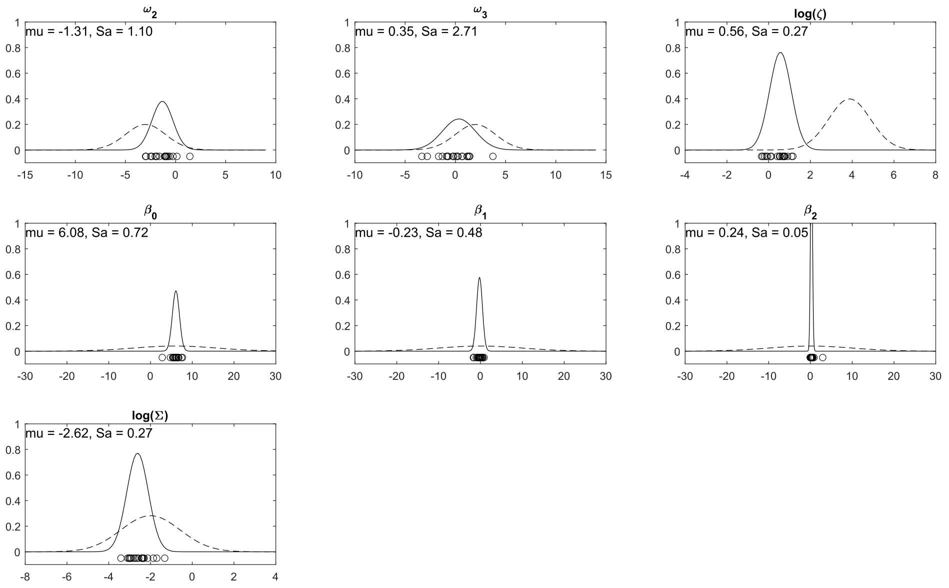


**Figure S2B** | *Empirical prior* densities of M2. For each free parameter of the model, the *initial prior* densities are shown as dashed line and the MAP estimates obtained from model inversion on the held-out data set ($N_{pilot}$) under the *initial priors* as black circles. The estimated *empirical priors* are shown as black line.

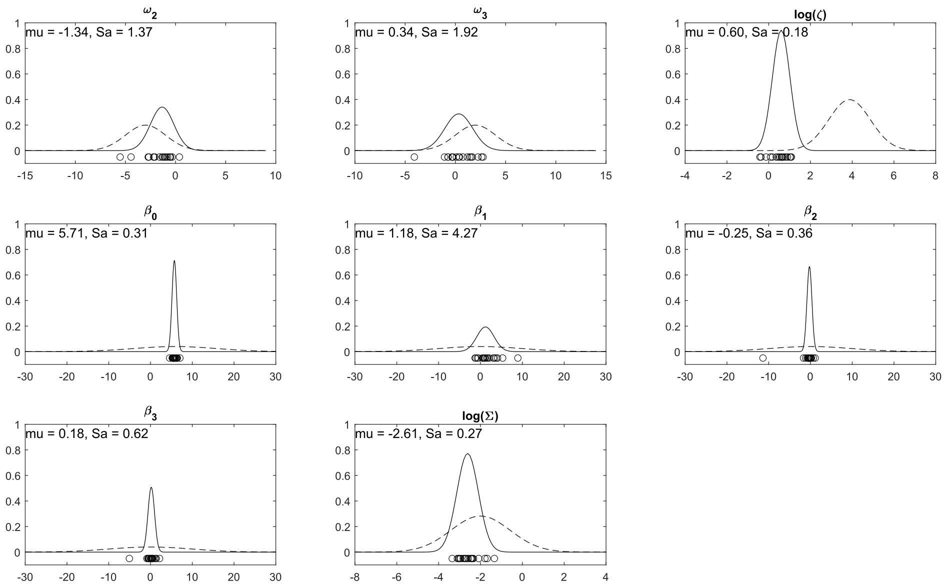


**Figure S2C** | *Empirical prior* densities of M3. For each free parameter of the model, the *initial prior* densities are shown as dashed line and the MAP estimates obtained from model inversion on the held-out data set ($N_{pilot}$) under the *initial priors* as black circles. The estimated *empirical priors* are shown as black line.

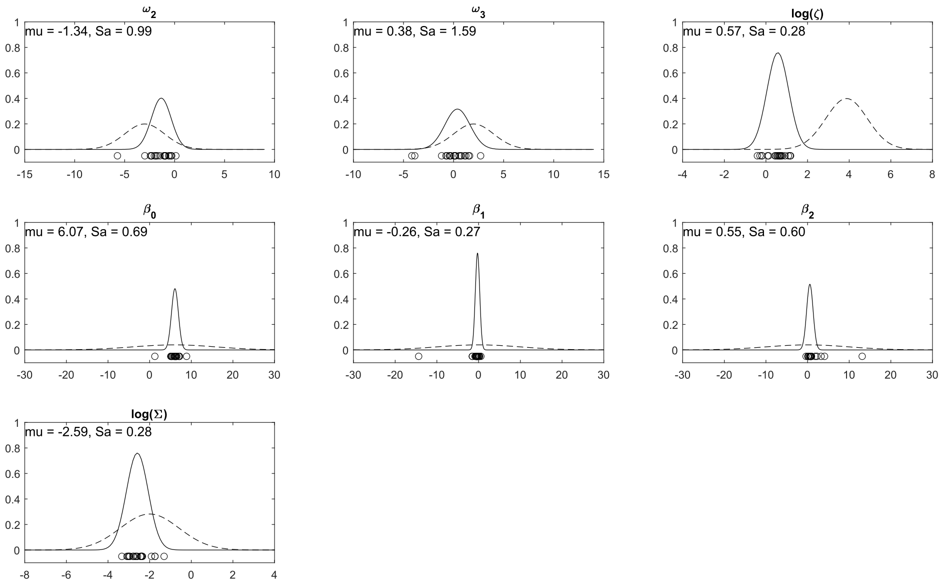


**Figure S2D** | *Empirical prior* densities of M4. For each free parameter of the model, the *initial prior* densities are shown as dashed line and the MAP estimates obtained from model inversion on the held-out data set ($N_{pilot}$) under the *initial priors* as black circles. The estimated *empirical priors* are shown as black line.

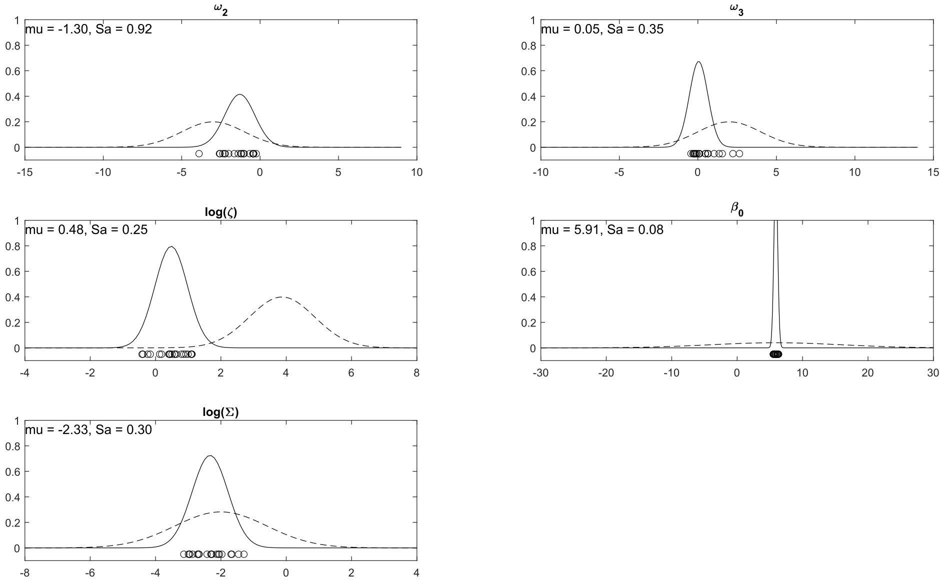


**Figure S2E** | *Empirical prior* densities of M5. For each free parameter of the model, the *initial prior* densities are shown as dashed line and the MAP estimates obtained from model inversion on the held-out data set ($N_{pilot}$) under the *initial priors* as black circles. The estimated *empirical priors* are shown as black line.

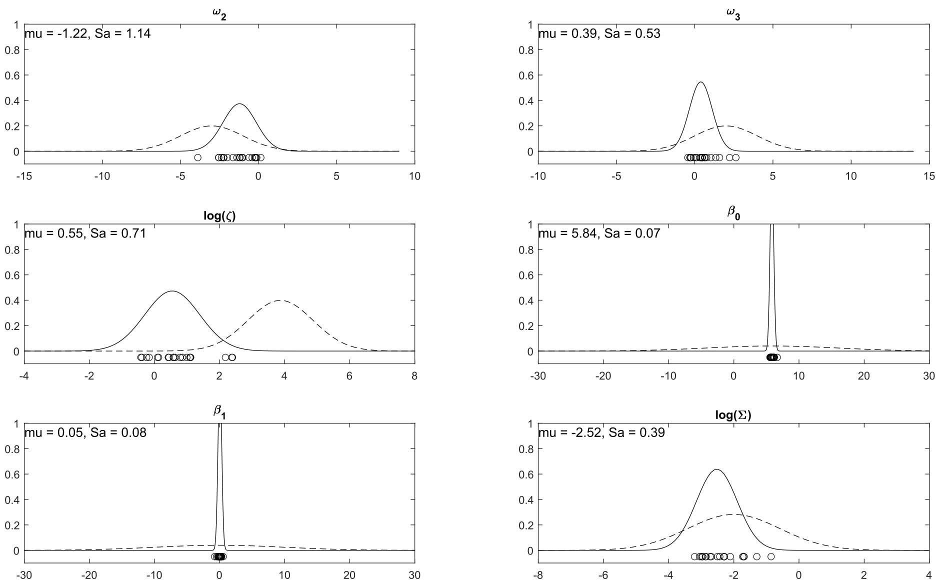


**Figure S2F** | *Empirical prior* densities of M6. For each free parameter of the model, the *initial prior* densities are shown as dashed line and the MAP estimates obtained from model inversion on the held-out data set ($N_{pilot}$) under the *initial priors* as black circles. The estimated *empirical priors* are shown as black line.

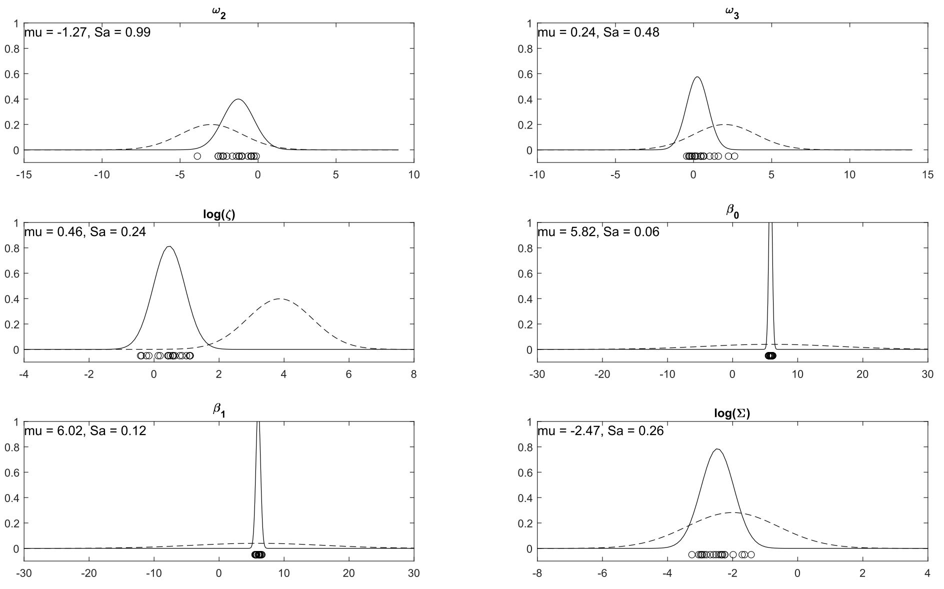


**Figure S2G** | *Empirical prior* densities of M7. For each free parameter of the model, the *initial prior* densities are shown as dashed line and the MAP estimates obtained from model inversion on the held-out data set ($N_{pilot}$) under the *initial priors* as black circles. The estimated *empirical priors* are shown as black line.

### S2b. *Empirical prior predictive distributions of M1-M7*

Empirical prior predictive distributions for the binary response modality and eHGF are shown in Figures S2H-S2N.

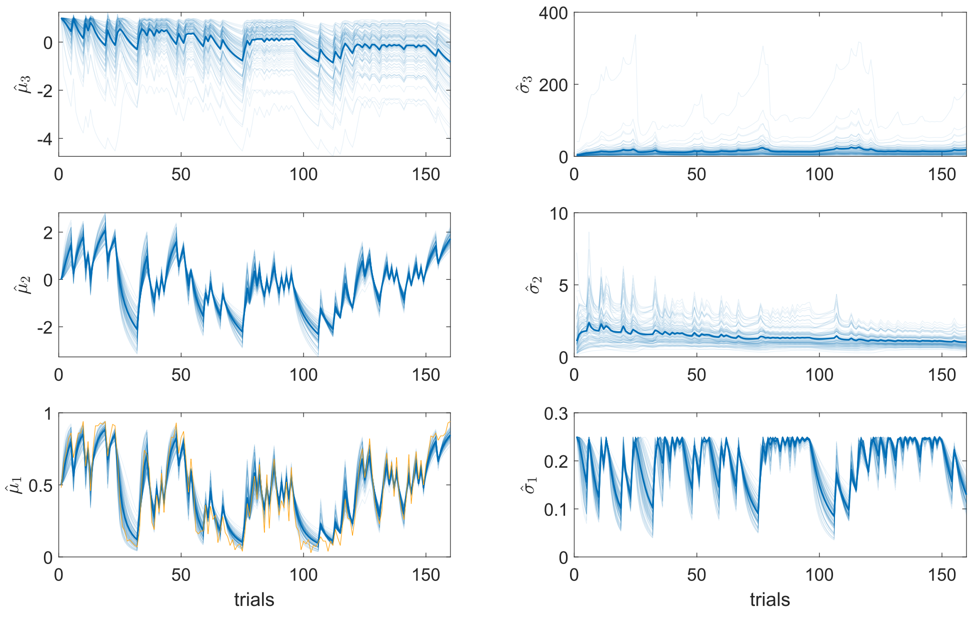


**Figure S2H** | *Empirical prior* predictive distribution of M1 (binary response modality and eHGF). Simulated belief trajectories ($N_{sim} = 100$) are shown for every level of the perceptual model. The mean belief is shown on the left whereas the uncertainty (variance) of the belief is shown on the right. The thick blue line represents the average over all simulated belief trajectories at every level. In the lower left panel, the yellow line represents the average simulated binary response.

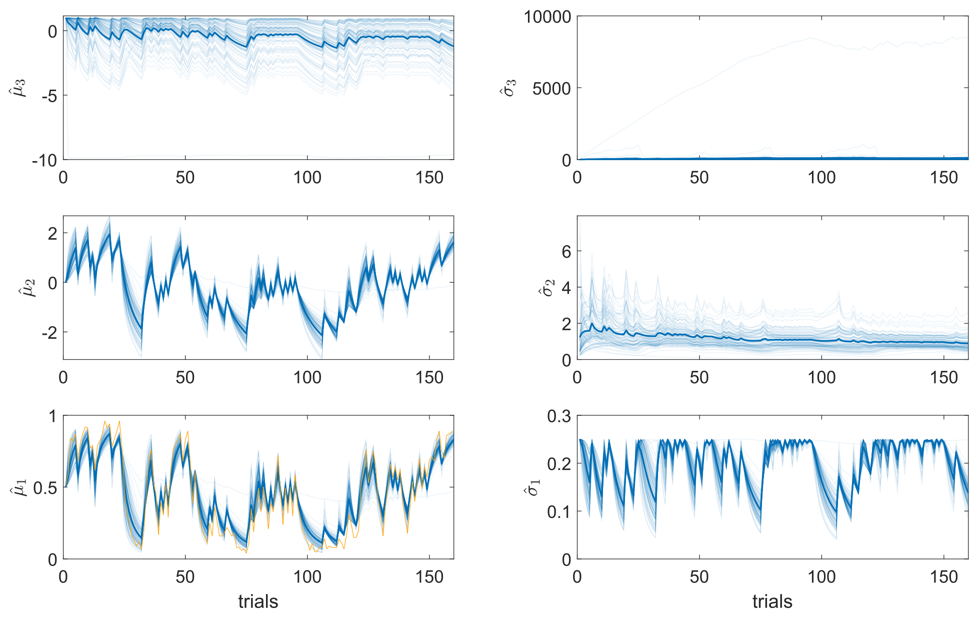


**Figure S2I** | *Empirical prior* predictive distribution of M2 (binary response modality and eHGF). Simulated belief trajectories ($N_{sim} = 100$) are shown for every level of the perceptual model. The mean belief is shown on the left whereas the uncertainty (variance) of the belief is shown on the right. The thick blue line represents the average over all simulated belief trajectories at every level. In the lower left panel, the yellow line represents the average simulated binary response.

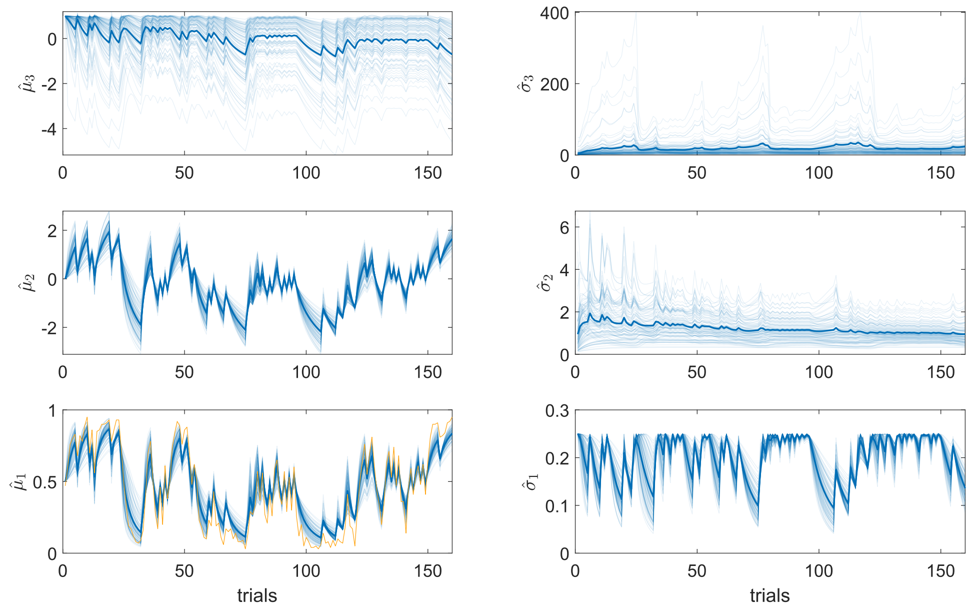


**Figure S2J** | *Empirical prior* predictive distribution of M3 (binary response modality and eHGF). Simulated belief trajectories ($N_{sim} = 100$) are shown for every level of the perceptual model. The mean belief is shown on the left whereas the uncertainty (variance) of the belief is shown on the right. The thick blue line represents the average over all simulated belief trajectories at every level. In the lower left panel, the yellow line represents the average simulated binary response.

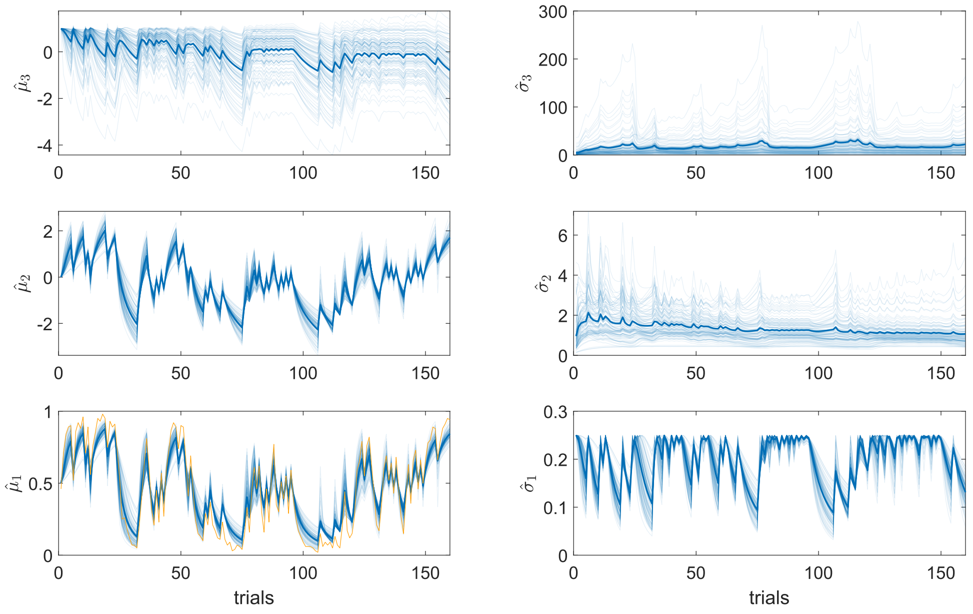


**Figure S2K** | *Empirical prior* predictive distribution of M4 (binary response modality and eHGF). Simulated belief trajectories ($N_{sim} = 100$) are shown for every level of the perceptual model. The mean belief is shown on the left whereas the uncertainty (variance) of the belief is shown on the right. The thick blue line represents the average over all simulated belief trajectories at every level. In the lower left panel, the yellow line represents the average simulated binary response.

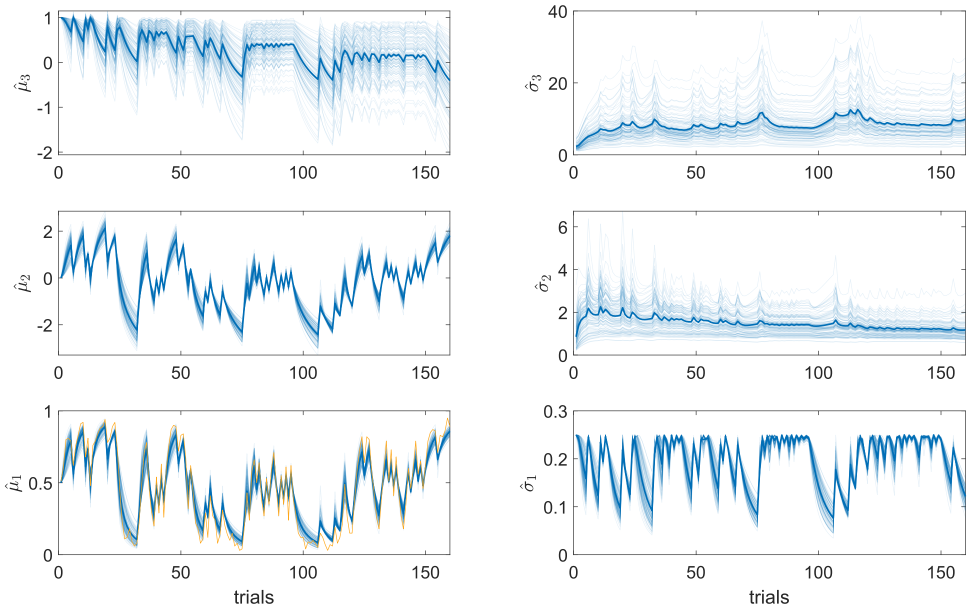


**Figure S2L** | *Empirical prior* predictive distribution of M5 (binary response modality and eHGF). Simulated belief trajectories ($N_{sim} = 100$) are shown for every level of the perceptual model. The mean belief is shown on the left whereas the uncertainty (variance) of the belief is shown on the right. The thick blue line represents the average over all simulated belief trajectories at every level. In the lower left panel, the yellow line represents the average simulated binary response.

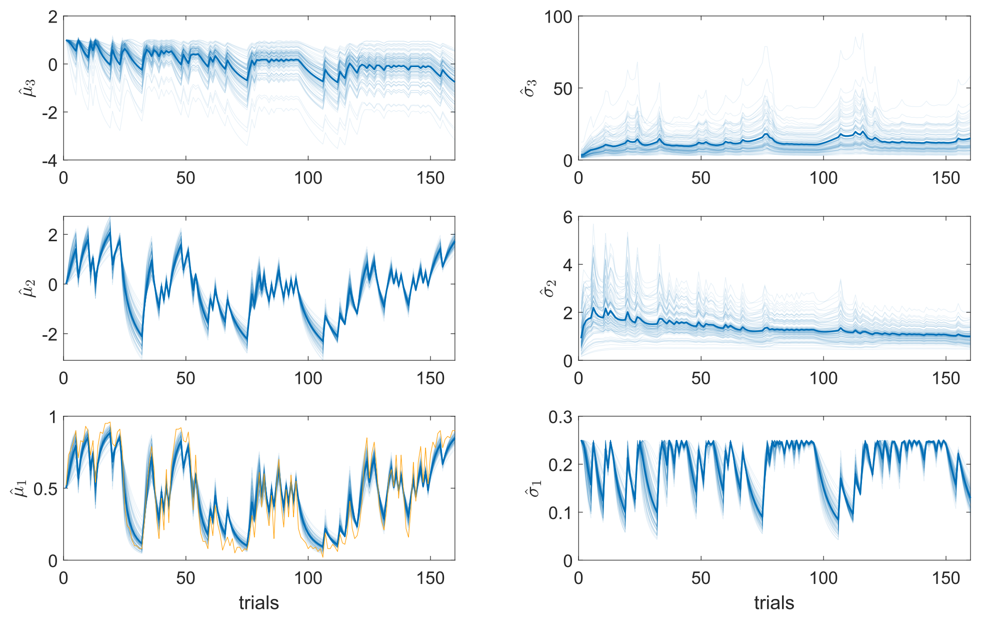


**Figure S2M** | *Empirical prior* predictive distribution of M6 (binary response modality and eHGF). Simulated belief trajectories ($N_{sim} = 100$) are shown for every level of the perceptual model. The mean belief is shown on the left whereas the uncertainty (variance) of the belief is shown on the right. The thick blue line represents the average over all simulated belief trajectories at every level. In the lower left panel, the yellow line represents the average simulated binary response.

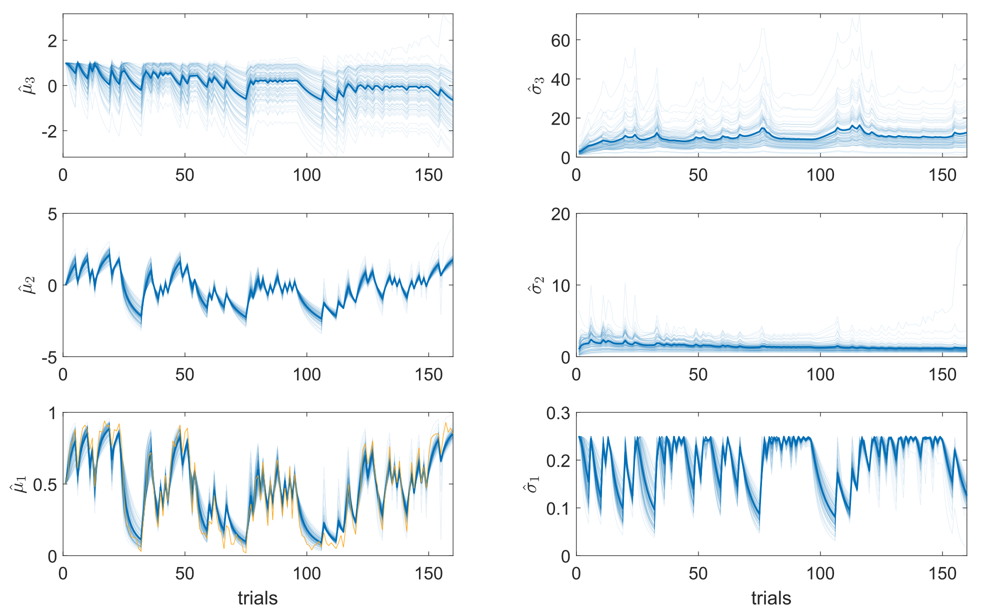


**Figure S2N** | *Empirical prior* predictive distribution of M7 (binary response modality and eHGF). Simulated belief trajectories ($N_{sim} = 100$) are shown for every level of the perceptual model. The mean belief is shown on the left whereas the uncertainty (variance) of the belief is shown on the right. The thick blue line represents the average over all simulated belief trajectories at every level. In the lower left panel, the yellow line represents the average simulated binary response.

Empirical prior predictive distributions for the continuous response modality are shown in Figures S2O and S2P.

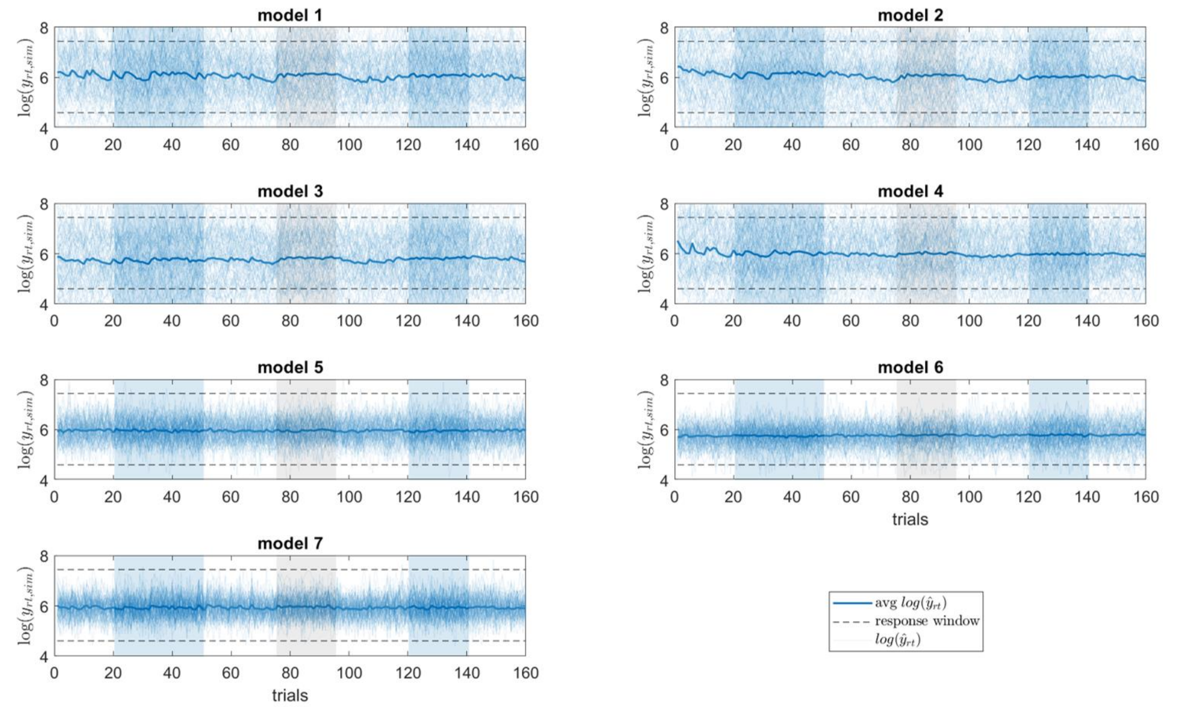


**Figure S2O** | Empirical prior predictive distributions of M1-M7 (simulated log RTs). Simulated log RT trajectories are shown in blue with the thick blue line representing the average over all simulated trajectories for each model ($N_{sim} = 100$). The dashed lines represent the boundaries of the response window in the SPIRL task.

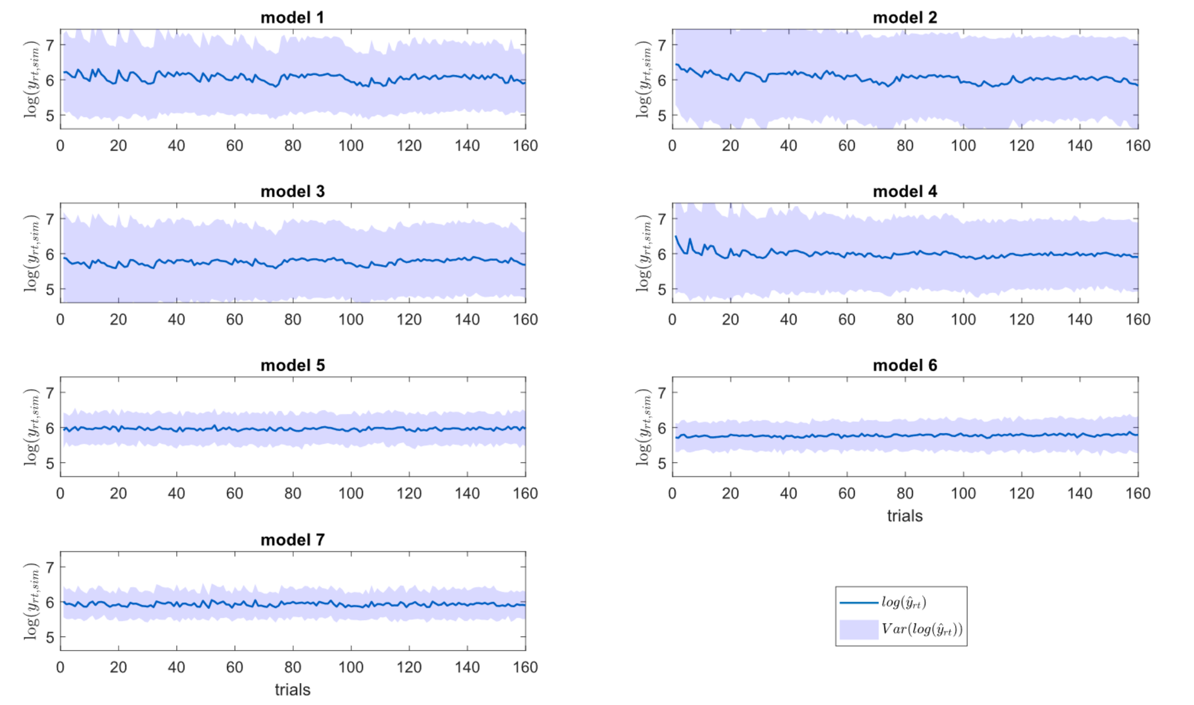


**Figure S2P** | Empirical prior predictive distributions of M1-M7 (simulated log RTs). Mean and standard deviation of simulated log RT trajectories for each model are shown in blue ($N_{sim} = 100$).

### S3: Parameter recovery results of M2-M7

Results from the parameter recovery analysis of M2-M7 are shown in Figures S3A-S3F.

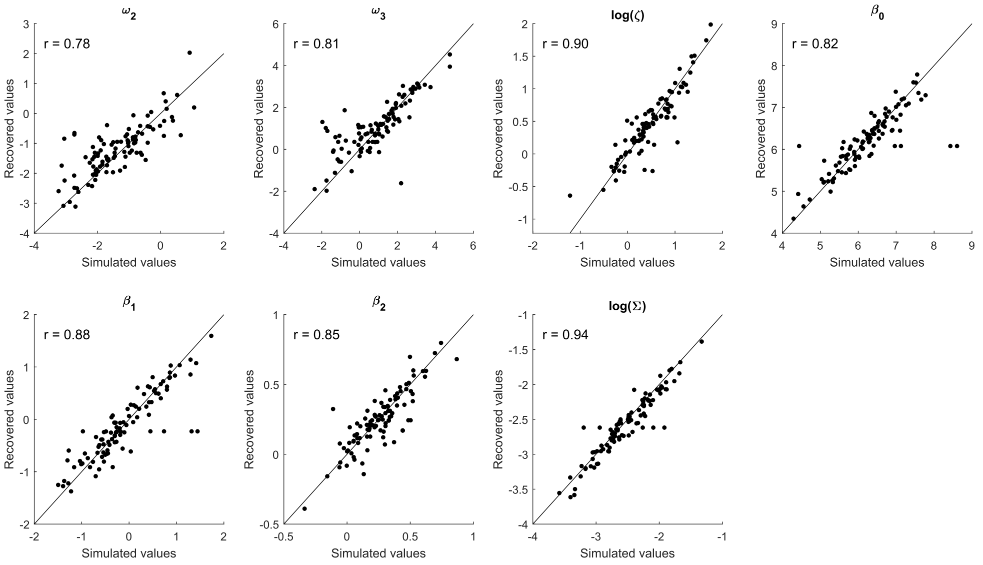


**Figure S3A** | Parameter recovery of M2. $N_{sim} = 100$ parameter values are displayed (simulated parameter values on the x-axis, fitted parameter values on the y-axis) and Pearson correlation coefficients between simulated and estimated parameter values for each free parameter are denoted by $r$. The black line is the identity line representing perfect recovery.

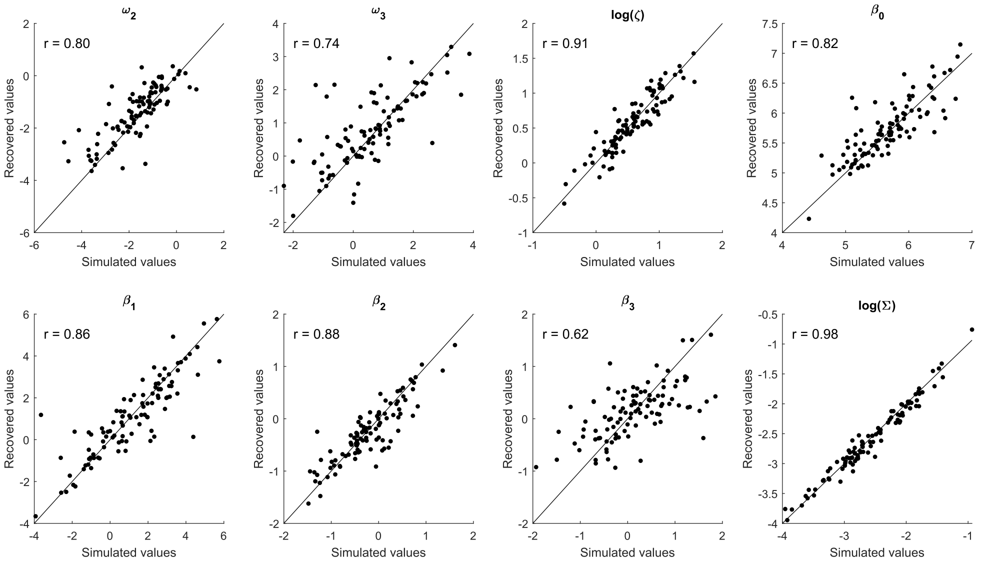


**Figure S3B** | Parameter recovery of M3. $N_{sim} = 100$ parameter values are displayed (simulated parameter values on the x-axis, fitted parameter values on the y-axis) and Pearson correlation coefficients between simulated and estimated parameter values for each free parameter

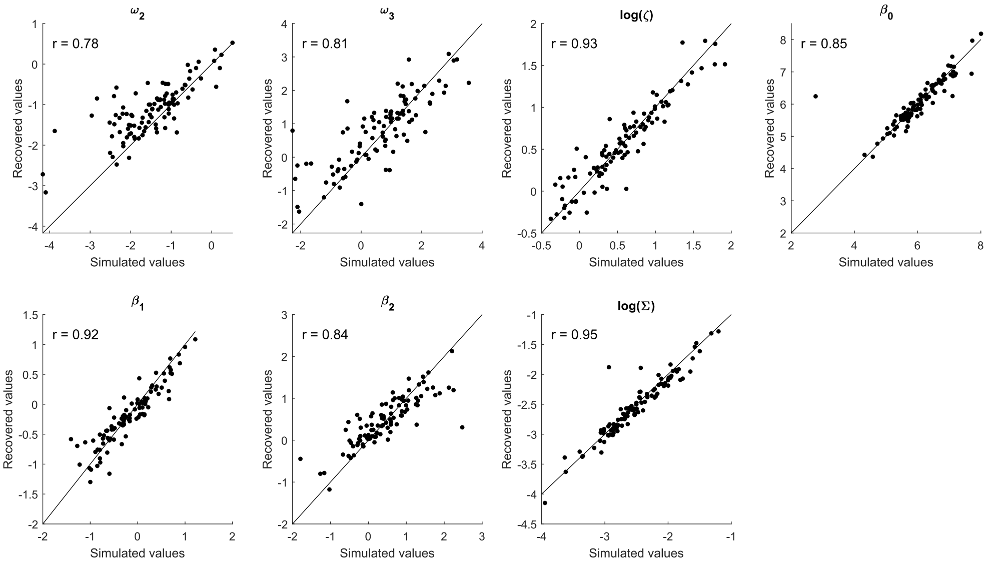


**Figure S3C** | Parameter recovery of M4. $N_{sim} = 100$ parameter values are displayed (simulated parameter values on the x-axis, fitted parameter values on the y-axis) and Pearson correlation coefficients between simulated and estimated parameter values for each free parameter

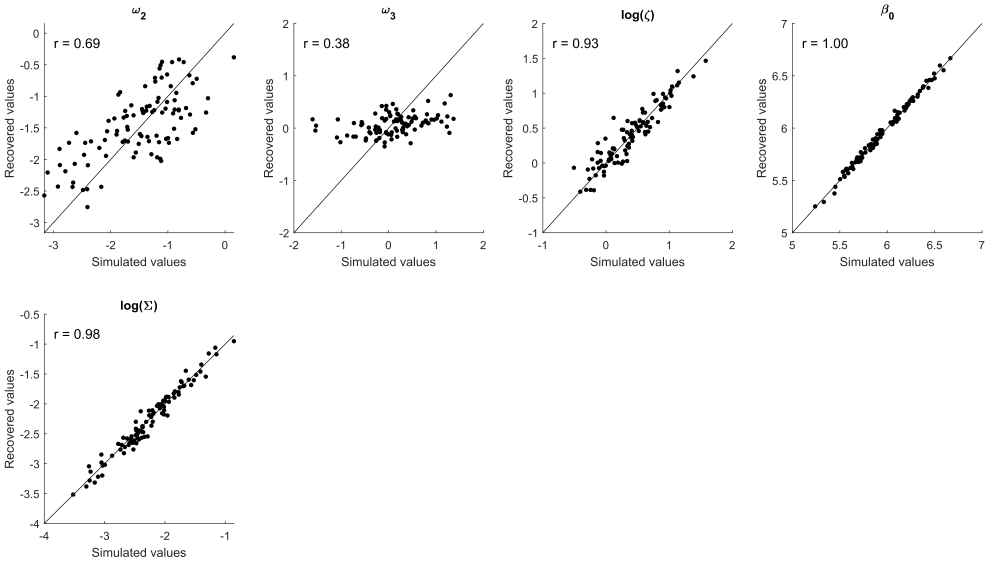


**Figure S3D** | Parameter recovery of M5. $N_{sim} = 100$ parameter values are displayed (simulated parameter values on the x-axis, fitted parameter values on the y-axis) and Pearson correlation coefficients between simulated and estimated parameter values for each free parameter

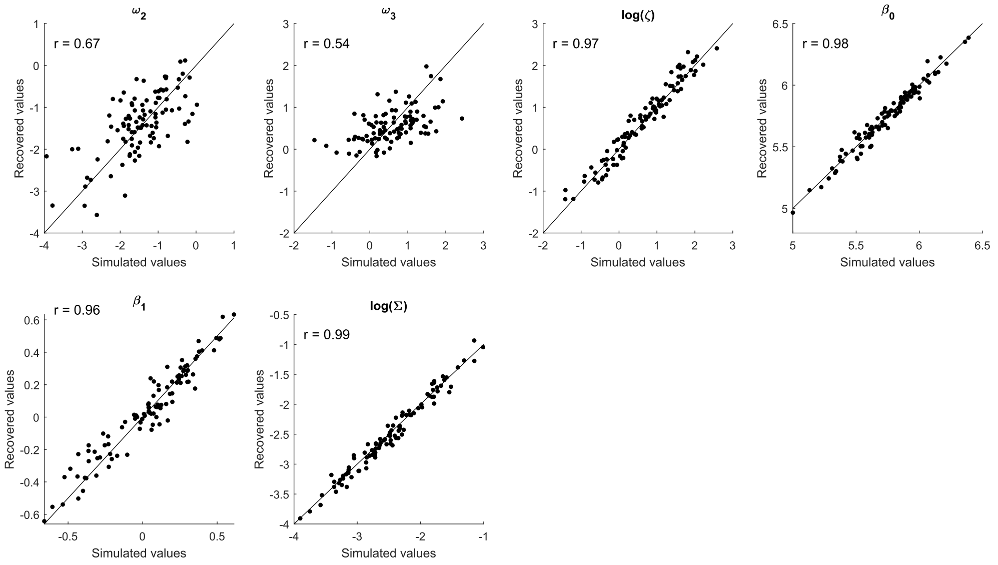


**Figure S3E** | Parameter recovery of M6. $N_{sim} = 100$ parameter values are displayed (simulated parameter values on the x-axis, fitted parameter values on the y-axis) and Pearson correlation coefficients between simulated and estimated parameter values for each free parameter

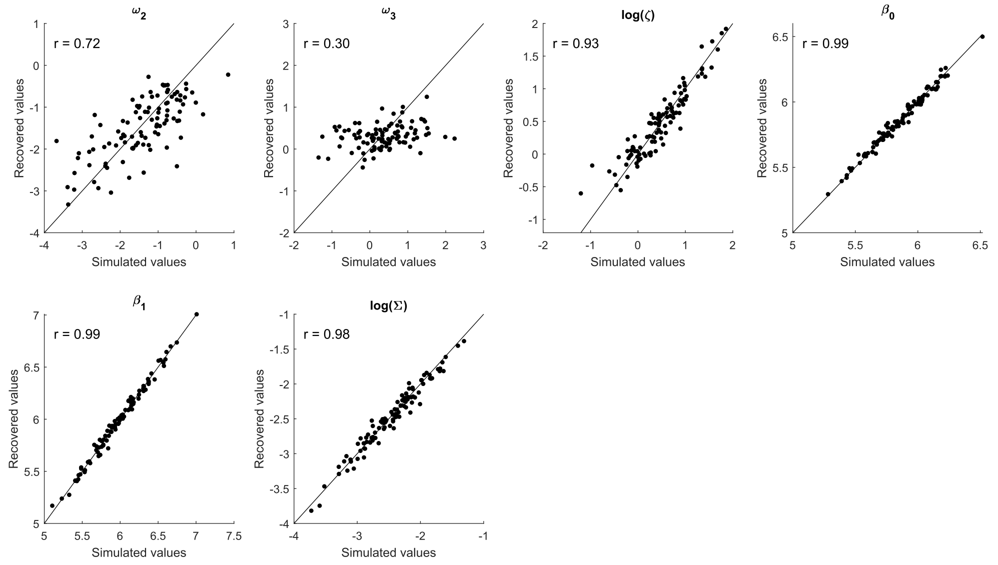


**Figure S3F** | Parameter recovery of M7. $N_{sim} = 100$ parameter values are displayed (simulated parameter values on the x-axis, fitted parameter values on the y-axis) and Pearson correlation coefficients between simulated and estimated parameter values for each free parameter

### S4: Average log RT trajectories and log RT model fits of M1-M7

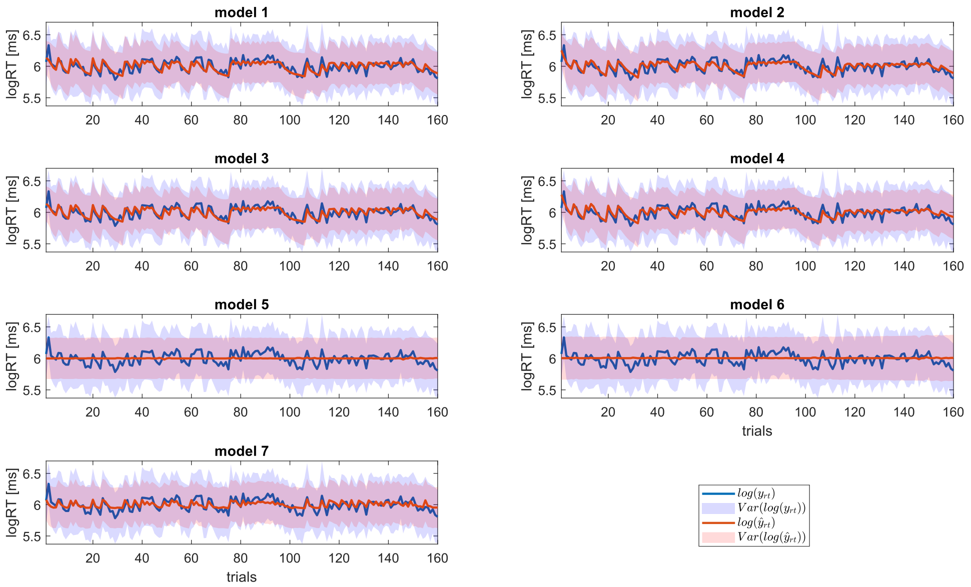


**Figure S4** | Average of the predicted (M1-M7) and average of the actual log RTs. In blue, mean and variance of log RT trajectories over all participants of the main data set are displayed ($N_{main} = 59$). In red, mean and variance of the log RT model fits is shown for each of the seven models in the model space.

### S5: Posterior predictive checks for M1

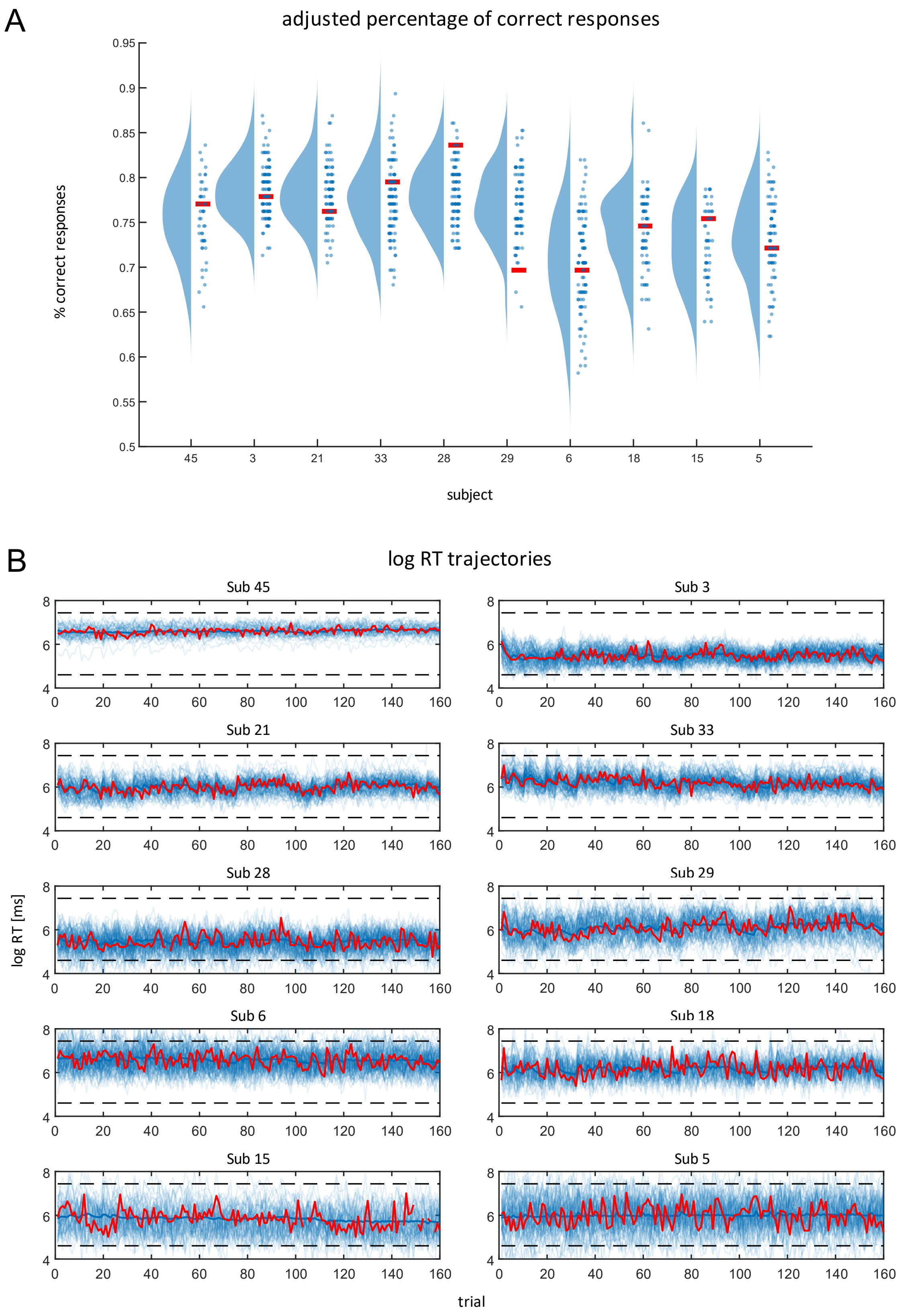


**Figure S5** | Posterior predictive checks for M1. Data from ten participants of the main data set are shown. Participants were chosen according to goodness of model fit of M1, i.e. the four participants with the highest log likelihood values (45, 3, 21, 33), two participants with average goodness of fit (28, 29) and the four participants showing the worst fit (6, 18, 15, 5). **A** displays adjusted correctness of binary responses for these participants in red. Blue circles are the simulated adjusted correctness values resulting from sampled parameter values of the subject-specific posteriors of M1. The blue probability densities are the estimated posterior predictive densities based on the samples drawn from the posteriors ($N_{ppc} = 100$) using kernel density estimation as implemented in the RainCloudPlots library. In **B**, we show empirical log RT trajectories of the ten participants in red. Fine blue lines are simulated log RT trajectories resulting from sampled parameter values of the subject-specific posteriors and the thick blue line represents the predicted log RT when using the MAP estimates of M1 for each participant to generate synthetic RT data.
